# Transient histone deacetylase inhibition reveals cell type invariant and specific effects of chromatin decondensation on irradiation response

**DOI:** 10.1101/2025.06.20.660786

**Authors:** Heng Li, Campbell Maben, Lindsey Young, Debjani Pal, Sandra Davern, Rachel Patton McCord

## Abstract

Radiation therapy plays a prominent role in breast cancer treatment, but the high doses of radiation damage both healthy and cancerous cells. Therefore, additional research is needed into combination therapies that could preferentially radiosensitize cancer cells compared to surrounding healthy tissue without causing deleterious side effects. Histone deacetylase inhibitor drugs (HDACis) have been tested as radiosensitizers in both basic research and clinical trials, but the long exposure time typically used in these treatments and the lack of matched healthy cell controls often leave aspects of their mechanism of action unclear. Here, we show that transient (2 hour) trichostatin A (TSA) treatment of cancerous and non-tumorigenic breast epithelial cell lines increases immediate DNA damage and decreases long term cell viability in both cell types at high radiation doses. Transient TSA treatment also causes an increase in DNA damage signals after 5 Gy in other cancer and healthy cell types: A375 melanoma cells and BJ5-ta fibroblasts. This suggests that chromatin decompaction acts to increase cellular vulnerability to initial DNA damage from high doses of radiation DNA damage in a cell type independent manner that does not rely on changes to DNA repair pathways caused by longer TSA treatment. However, responses to lower doses of radiation and long term survival are more cell type specific: only MCF7 cells experience an effect of TSA on DNA damage after 1 Gy radiation while MCF10a cells experience somewhat more evident cell viability effects of combined TSA and radiation treatment long term.

**Scope statement:** This manuscript covers several topics that are core to the mission of Frontiers in Cell and Developmental Biology, including cancer cell biology as compared to non-cancerous cell function, cell death vs. survival, and epigenetics and chromosome structure. While some of the implications of this work touch on cancer therapeutics, the study itself focuses on the basics of the interplay between genome architecture and DNA damage and cellular survival / response. We noted several recent articles on related topics in this journal, including the effects of combination treatments on cancer cells (doi 10.3389/fcell.2025.1636288), radiotherapy effects on cells (doi 10.3389/fcell.2025.1568634), and DNA damage (doi 10.3389/fcell.2025.1575483). Our work is well suited for a Brief Research Report as it provides key but focused data on comparisons of radiosensitivity in response to transient chromatin decompaction in cancer vs. healthy cells.

## Introduction

Radiotherapy is a primary treatment for cancers such as breast cancer, but high-energy radiation can damage DNA in both cancer cells and surrounding healthy tissues. Therefore, extensive research has sought to preferentially radiosensitize cancer cells compared to healthy cells and to understand how different cell types respond to radiation (Citrin and Mitchell, 2014). Previous work suggests that radiation-induced DNA damage can be impacted by chromatin compaction and histone modification state. Decondensed nuclei showed more DNA damage signal upon high-dose irradiation than condensed nuclei (Takata et al., 2013). A computational simulation similarly predicted that DNA compaction influences radiation-induced DNA damage (Tang et al., 2019). Other imaging evidence has shown that ionizing radiation results in DNA repair foci preferentially localized in less compact euchromatin. (Cowell et al., 2007, Falk et al., 2008) Indeed, DNA occupancy by histone proteins can absorb the effect of radiation and protect DNA from damage (Warters and Lyons, 1992, Nygren et al., 1995, Brambilla et al., 2020). Therefore, chromatin compaction is a potential target for radiosensitivity.

Generally, heterochromatic regions tend to be methylated on histone H3 at lysine 9 and/or lysine 27, while euchromatic regions instead exhibit acetylation on these same histone tails. (Lachner et al., 2001, Yan et al., 2018, Cao et al., 2002, Ferrari et al., 2014). Histone acetylation neutralizes the positive charges on lysine-rich histone tails, weakening their interactions with the negatively charged DNA and making chromatin less compact and more accessible (Kim et al., 2023). Trichostatin A (TSA) is a pan-Histone deacetylase inhibitor (HDACi), that results in increases in histone acetylation, chromatin decompaction (Wang et al., 2009), and chromosome structure changes (Stephens et al., 2018, Vinayak et al., 2025). TSA has been previously studied as an anti-cancer agent (Bouyahya et al., 2022) and was found to increase the radiosensitivity of numerous types of human cancer cells, such as human colon cancer, bladder cancer, non-small-cell lung cancer, chronic myelogenous leukemia, squamous cell carcinoma, and breast cancer (Kim et al., 2010, Zhang et al., 2009, Yu et al., 2012, He et al., 2014, Jia et al., 2017, Sun et al., 2014, Karagiannis et al., 2005, Paillas et al., 2020). These studies have shown that long-term treatments (12 h – 96 h) with TSA can sensitize cancerous cells to radiation by inducing cell cycle arrest, diminishing DNA damage repair, inhibiting cell autophagy, and enhancing cell apoptosis. However, the numerous secondary effects and impacts on cell viability from TSA alone that happen with long-term treatment make it difficult to assess the contribution of chromatin decompaction itself to the amount of DNA damage experienced by cancer cells and the radiosensitizing effects of TSA treatment. Indeed, previous literature has demonstrated substantial cytotoxicity, reduced proliferation, and cell cycle arrest effects of even 100 nM TSA treatment alone on breast cancer cell lines if the treatment is applied for 24 h or more (Alao et al., 2004, Min et al., 2004, Kong et al., 2019, Noh et al., 2016, Urbinati et al., 2011, Vigushin et al., 2001).

Such a long TSA treatment is not needed to produce effects on chromatin acetylation and chromatin compaction. The increases in histone acetylation caused by TSA happen rapidly, with substantial changes even detected within 30 minutes (Wang et al., 2009), so a much shorter treatment than 12 hours could be used to evaluate the effect of chromatin changes on radiosensitivity more directly. In other contexts, transient TSA treatments (3 hours) induced chromatin decondensation and nuclear deformation in adult meniscal fibrochondrocytes (Heo et al., 2020). Likewise, our previous studies indicated that 500 nM TSA treatment for 2 hours was sufficient to increase histone acetylation and chromatin accessibility and change chromosome structure in A375 cells (Das et al., 2020, Vinayak et al., 2025). Therefore, here, we aim to test the hypothesis that increasing chromatin accessibility by transient (2 hour) TSA treatment will enhance cell sensitivity to radiation.

Another key factor often missing from existing HDACi radiosensitivity studies is a matched comparison between healthy and cancer cells. Many studies demonstrate the radiosensitizing effects of TSA on cancer cells, but with no comparison of healthy cell response, it is unclear whether the treatment would influence all cell types equally or preferentially target cancer cells. Here, we use non-tumorigenic breast epithelial cells (MCF-10A) and human breast cancer cells (MCF-7) to investigate the influence of initial genome structure on the cellular response to X-ray radiation. Unlike other breast cancer cell lines (BT-474 and MDA-MB-231), MCF-7 cells do not exhibit pre-existing mutations in DNA damage repair pathways (Gingrich et al., 2025). We find that 2 hours of TSA treatment increases the acetylation level in both healthy and cancerous breast epithelial cells. This transient TSA treatment increases the DNA damage caused by subsequent 5 Gy irradiation in both MCF-10A and MCF-7 cells. However, only MCF-7 cells, not MCF-10A cells, show increased DNA damage after TSA treatment followed by a lower dose of irradiation (1 Gy). These different responses to radiation after TSA treatment led to a distinction in long-term cell survival rate, indicating that transient pre-treatment of TSA enhances the radiosensitivity to breast cancer cells, but not normal breast cells, to moderate doses of radiation.

## Materials and Methods

### Cell culture

MCF-10A (RRID: CVCL_0598) and MCF-7 (RRID:CVCL_0031) cells were purchased from ATCC (CRL-10317 and HTB-22 respectively). MCF-10A cells were cultured in Mammary Epithelial Cell Growth Medium BulletKit (Lonza CC-3150; following the ATCC instruction, 100 ng/ml Cholera toxin was used to make the complete growth medium instead of GA-1000 provided with kit) at 37°C supplied with 5% CO_2_. MCF-7 cells were cultured in complete Eagle’s Minimum Essential Medium (ATCC 30-2003; 10% FBS. 1% Pen-Strep, 10 µg/ml Insulin) at 37°C supplied with 5% CO_2_. A375 cells (RRID:CVCL_0132) were purchased from ATCC (CRL-1619) and cultured in complete DMEM medium (Corning-10-013-CV; 10% FBS, 1% Pen-Strep, 1% L-Glutamine) at 37° C supplied with 5% CO2. BJ-5ta cells were purchased from ATCC (CRL-4001) and cultured in a 4:1 ratio of DMEM (Corning, 10-013-CV) and 1x Medium 199 (Gibco, 11-150-059), supplemented with 10% FBS (Corning, 35-010-CV), 0.01 mg/mL hygromycin B (Corning, 30-240-CR), 1% Pen-Strep (Gibco, 14140-122), and 1% L-Glutamate (Gibco, 25030-081). Cells were verified to be negative for mycoplasma.

### HDACi treatment

TSA was purchased from Tocris (1406, UK), dissolved in dimethyl sulfoxide (4-X, ATCC) and stored at -20°C with a stock concentration of 10 mM. For all treatments, TSA was diluted in cell culture media to the indicated concentrations and then cells were returned to the incubator for 2 hours. TSA was then washed off and fresh media added immediately prior to subsequent cell viability, irradiation, and acetylation assays.

### X-ray irradiation

Cells were irradiated using a RS 2000 X-ray Irradiator (RadSource). A 2 Gy/min dose rate was used to achieve the indicated total doses (160 kV; 25 mA). After irradiation, cells were transferred back to an incubator and kept at 37°C with 5% CO_2_.

### Cell proliferation and viability assays

#### MTS assay

Cell viability was determined using the CellTiter 96 Aqueous One Solution Cell Proliferation Assay System (Promega) with novel tetrazolium compound [3-(4,5-dimethylthiazol-2-yl)-5-(3-carboxymethoxyphenyl)-2-(4-sulfophenyl)-2H-tetrazolium, inner salt; MTS]. To determine the optimal incubation time for cellular reduction of MTS to produce soluble formazan, various numbers of MCF-10A and MCF-7 cells were added to 96 well plates and incubated with MTS reagent for 1-4 hours. The linear regression analysis showed the coefficient of determination (r^2^) was above 0.995 in both MCF-10A and MCF-7 cells under 2 hours incubation, indicating a linear response between cell number and absorbance of formazan product at 490 nm (Supplementary Fig. S1). For treatment assays, MCF-10A and MCF-7 cells were seeded in 96-well plates, at an initial density of 25,000 cells per 1 ml of medium, 24 h before TSA treatments. Cells were incubated with various concentrations of TSA (250 nM, 500 nM, 750 nM, 1000 nM) for 2 h; cells with no TSA (DMSO only) were included for the negative control. After TSA treatment, 20µL MTS reagent was added into each well and incubated at 37°C with humidified 5% CO_2_ for 2 h. Absorbance at 490 nm was measured using a microplate reader (BioTek 800 TS, Agilent).

#### Trypan blue exclusion assay

Cell proliferation and viability were measured using the trypan blue exclusion assay. MCF-10A and MCF-7 cells were seeded in 6-well plates, at an initial density of 1,000,000 cells per 1 mL of medium, 24 h before TSA treatments, respectively. Cells were incubated with various concentrations of TSA (0 nM, 250 nM, 500 nM, 750 nM, 1000 nM) for 2 h; cells with fresh medium were included for the negative control. After TSA treatment, cells were trypsinized and centrifuged at 1,000 x g for 5 minutes. An aliquot of cell suspension was mixed with 0.4% trypan blue and loaded on a hemacytometer. Cell counts were examined using the automated cell counter (Corning).

#### Immunofluorescence

MCF-10A, MCF-7, BJ5ta, or A375 cells were seeded on Poly-D-Lysine coated 35 mm dishes with No. 1.5 glass coverslips (MatTek) prior to TSA treatment and X-ray exposure. Cells were treated with DMSO or TSA (500 nM) for 2 h. For X-ray exposure, TSA was washed out and replaced with fresh standard culture medium and then cells were exposed to the indicated dose of X-ray. Cells were fixed in 4% formaldehyde for 20 minutes at room temperature and washed three times with PBS for 5 minutes each. Cells were permeabilized and blocked in blocking buffer (10% goat serum, 0.5% triton in PBS) for 1h at room temperature, following which the primary antibody H3K9ac (RRID:AB_732920, Abcam, ab32129, 1:250), γ-H2AX (RRID:AB_2799949, Cell Signaling Technology, CST-80312, 1:250) was diluted into antibody dilution buffer (5% goat serum, 0.25% triton in PBS) and incubated overnight at 4°C. Cells were washed in PBS for 5 minutes three times and incubated in secondary antibody buffer (Alexa fluor 488 goat antibody anti-mouse, RRID:AB_2556548, Invitrogen, R37120) for 30 minutes at room temperature in the dark. Cells were then washed with PBS for 5 minutes three times. Coverslips were removed and put on slides with ProLong Diamond Antifade Mountant with DAPI (Invitrogen, P36962). Samples were stored at 4°C overnight and imaged using a Leica SP8 confocal microscope equipped with a 63x oil immersion objective.

#### Western Blot

After TSA treatment with indicated concentrations, cells were trypsinized and collected by centrifugation at 1,000 x g for 5 minutes. Cells were resuspended in 300 ml of RIPA buffer (Thermo Scientific, I89900), supplemented with protease inhibitors/EDTA (GenDepot 50-101-5485), and phosphatase inhibitors (GenDepot 50-101-5488). Cells were lysed by repeated pipetting followed by incubation, on ice, for 10 min. DNA was degraded with micrococcal nuclease (Thermo Scientific, FEREN0181). Cell debris was removed by centrifugation at 14,000 rcf for 15 min. Protein samples were quantitated with the BCA Protein Assay Kit (Thermo 234225) and stored at −80 °C until use.

Denatured protein samples were resolved using 4–12% Bis-Tris Plus gels (Invitrogen, NW04120BOX). Protein was transferred to low fluorescence PVDF membranes using the mini Bolt Blotting System (Invitrogen, B1000). Blotting was performed using the antibodies: GAPDH (RRID:AB_307275, Abcam, ab9485, 1:2500) and acetylated histone H3 (RRID:AB_2115283, Millipore, 06-599, 1:1000). Detection was carried out using the secondary antibody: goat anti-rabbit (1:10,000; IRDye 680RD [92568071]; Licor). Images of fluorescent signal were acquired using an Odyssey DLx Imager and quantitated using the Image Studio software (LICOR). All uncropped blot images are presented in Supplementary Fig. S3.

#### Clonogenic assay

The clonogenic assay was used to determine cell survival after X-ray exposure and adapted as described previously(De et al., 2019). MCF-10A and MCF-7 cells were seeded at an initial density of 1,000 cells/well and 1,500 cells/well in 6-well plates, respectively and allowed to attach for 24 h. Cells were treated with DMSO or TSA (500 nM) for 2 h. After the TSA was washed out and replaced with fresh standard culture medium, cells were exposed to the indicated dose (0, 1, 2, or 5 Gy) of X-rays. Cells were then transferred back to an incubator and kept at 37°C with 5% CO_2_ for 14 days. Colonies were fixed and stained with 0.5% crystal violet containing 10% methanol prepared in PBS. Images were acquired using the Odyssey DLx Imaging System (LICORbio). Colonies were counted manually under the microscope by observing the size of colonies containing at least 50 cells and then counting the number of colonies that exceeded this threshold (Franken et al., 2006). Due to difficulties with interpreting plating efficiency based on colony counts for cells with spreading growth patterns like MCF-10A (Brix et al., 2020), as an alternative quantification method, ImageJ was used on captured images to calculate the percent area covered by colonies. This was accomplished using the published ColonyArea ImageJ plugin (Guzmán et al., 2014) available at https://imagej.net/plugins/colonyarea. Survival curves were fit with a linear quadratic equation using GraphPad Prism version 10.6.1 and sensitizer enhancement ratios were calculated at survival fraction 50% as described in (Kim et al., 2010).

#### ImageJ analysis

Confocal microscopy images (.lif) were loaded into ImageJ (RRID:SCR_003070) with the Bio-Formats Importer plugin. For each image, the slices of the Z-stack were summed. Nuclei boundaries were established through thresholding of the DAPI signal. Merged nuclei were split with the Adjustable Watershed plugin. Nuclei with areas below 50 μm^2^ were excluded. To measure signal intensity, the mean grey value of γH2AX or H3K9ac signal across Z-slices was recorded for each bounded nucleus. Nuclei were excluded from quantification if the γH2AX signal was widespread and uniform rather than in foci. Foci counting quantification was also performed on 1 Gy irradiation conditions for MCF-7 and MCF-10A cells. γH2AX foci were automatically counted with the count_points plugin, with nuclei at the edge or nuclei that were non-singular excluded from the analysis. As previously described, foci counting is not possible at higher irradiation doses as the damage becomes too dense for individual foci to be detected (Kurashige et al., 2016). For A375 cells, high levels of endogenous DNA damage made γH2AX signal quantification more challenging. To focus on the additional damage caused by irradiation, a threshold was chosen that isolated only bright foci in non-irradiated cells and then all images were subjected to this same threshold before analysis. To match A375 analysis, BJ5-ta cells were analyzed in the same manner using thresholding.

### Data availability

Confocal images and other relevant raw data are available at Dryad: DOI: 10.5061/dryad.bvq83bkr6

## Results

### Transient (two hour) TSA treatment does not alter cell viability or proliferation in healthy and cancerous breast epithelial cells

To assess the impact of chromatin decompaction on cancer and healthy cell radiosensitivity, we used 2-hour TSA treatment of non-cancerous MCF-10A breast epithelial cells and breast cancer MCF-7 cells. This length of treatment has been shown to change acetylation and chromatin compaction in various cell types in other studies (Wang et al., 2009, Vinayak et al., 2025). To determine how this TSA treatment affects the viability of MCF-10A and MCF-7 cell lines, we examined the cell metabolic activity with the MTS assay (**Supplementary Fig S1**). The cell viability was not affected by this transient TSA treatment when the concentration of TSA was increased up to 1000 nM in both healthy and cancer cells (**Fig. 1a**). Further, a trypan blue exclusion assay showed no significant differences in cell proliferation or cell death after 250-1000 nM TSA treatment in either MCF-10A or MCF-7 cells (**Fig. 1b**). These results are in agreement with other recent work which has shown that brief treatments with TSA can cause substantial changes to histone acetylation without dramatically affecting the rest of the acetylome or the cell cycle (Paldi et al., 2026).

**Figure 1.**
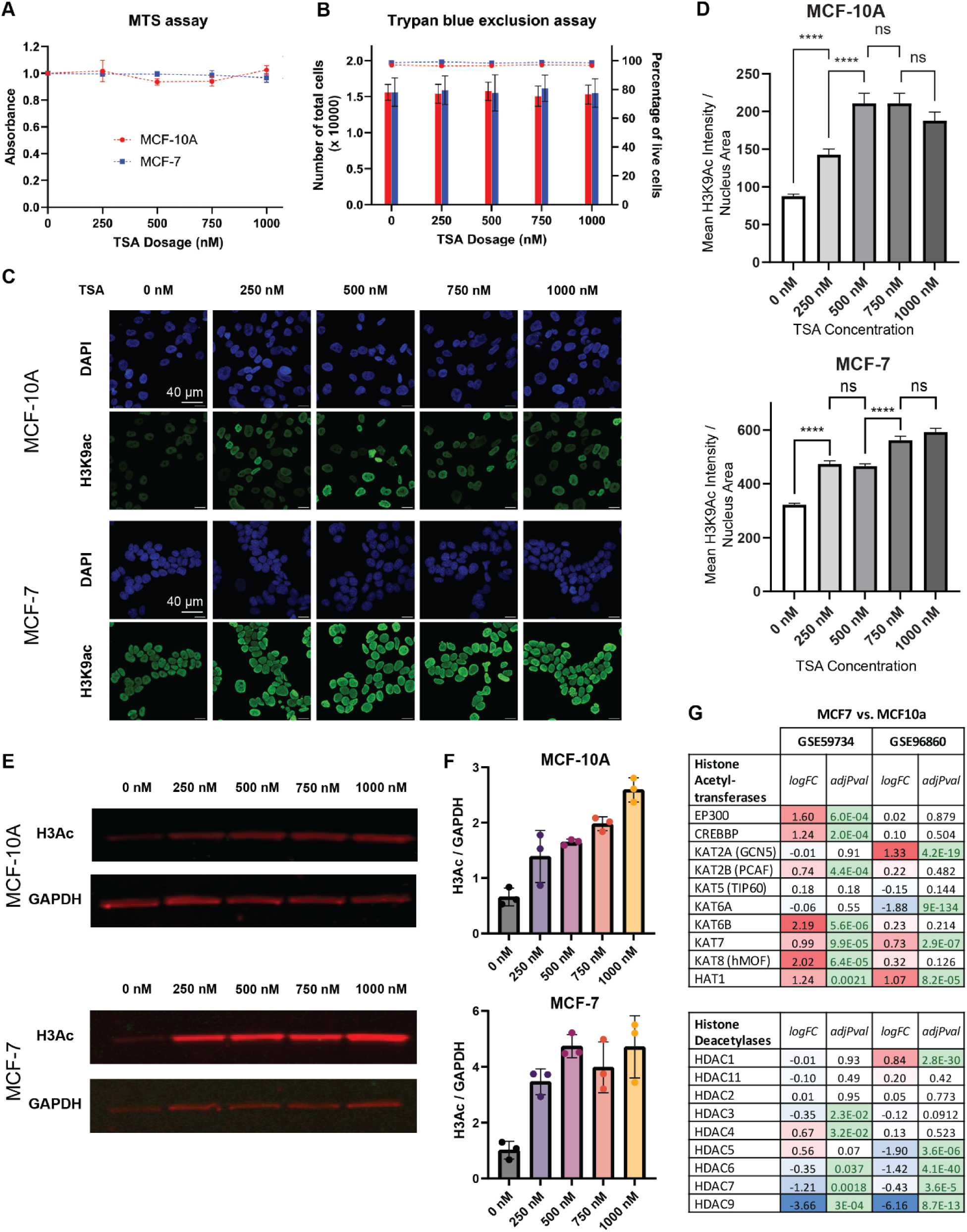
Two hour TSA treatment does not impact cell viability but increases histone acetylation in cancer and non-cancerous cells. **(A)** Influence of 2 hr TSA treatment (0, 250, 500, 750 and 1000 nM) on metabolic activity of MCF-10A (red) and MCF-7 (blue) cells, using the MTS assay. Data is presented as mean ± SEM of 3 biological replicates which each contain 6 technical replicates. MTS absorbance values are normalized to the value of the no treatment (0 nM) well. **(B)** Influence of 2 hr TSA treatment (0, 250, 500, 750 and 1000 nM) on cell viability of MCF-10A (red) and MCF-7 (blue) cells, using a trypan blue exclusion assay. Bars indicate the mean ± SEM of total cell counts for 3 biological replicates (left axis). Points along the dotted line indicate the percentage of live cells (negative for trypan blue) in each condition (right axis). **(C)** MCF-10A (top) and MCF-7 (bottom) cells were treated with indicated concentration of TSA (0, 250, 500, 750 and 1000 nM) for 2 h, then stained with DAPI (Blue) and anti-H3K9ac (Green). Images are maximum projections of Z-stacks. Scale bar: 40 µm (shown in top left image, same for all other panels). **(D)** Quantification of average H3K9ac intensity per nucleus. Bars show mean +/- SEM averaged across all measured nuclei (5 fields imaged per condition, N=150 – 300 nuclei per condition) (**** p < 0.0001 for pairwise one-tailed t-tests between each increasing concentration) **(E)** Western blot for total acetylated Histone H3 levels (H3Ac) compared to GAPDH loading control for MCF-10A and MCF-7 after indicated doses of 2 hr TSA treatment. Additional replicates and uncropped blots shown in Supplementary Figure S3. **(F)** Quantification of Western blot results across all 3 replicates. Bars indicate the mean ± SD of acetylated Histone H3 normalized by GAPDH loading control for 3 biological replicates (also shown as individual points). **(G)** Analysis of two publicly available RNA-seq datasets shows upregulation of histone acetyltransferases and downregulation of HDACs in MCF-7 vs. MCF-10A (in untreated conditions). logFC = log2(MCF-7/MCF-10A) RNA expression calculated by GEO2R. adjPval = False Discovery Rate adjusted p value for each comparison. Blue/Red colorscale indicates whether genes are expressed at lower (blue) or higher (red) levels in MCF-7 compared to MCF-10A. Green highlights comparisons with <5% FDR.

### Transient TSA treatment increases histone acetylation in both healthy and cancerous breast epithelial cells

To quantify the increase of acetylation caused by transient TSA treatment, we measured the immunofluorescence signal of H3K9Ac and performed a Western blot for total H3Ac. In both healthy cells (MCF-10A) and cancerous cells (MCF-7), we observed an elevated level of acetylation of H3K9 in cells exposed to TSA for 2 hours **(Fig. 1c-d, Supplementary Fig. S2)**. Total H3 acetylation also increased with increasing TSA concentration in both cell lines (**Fig. 1e, Supplementary Fig. S3**). Comparing the quantification of immunofluorescence and western blot signals, we observed that acetylation signal reached its maximum between 250 and 750 nM treatment concentrations, so we selected the 500 nM concentration as a midpoint for downstream radiosensitivity testing (**Fig. 1d, f**). Both immunofluorescence and western blot results showed higher signal intensity for acetylation after TSA treatment in MCF-7 compared to MCF-10A cells. This agrees with previous results showing that 48 h TSA treatment across this same concentration range produces higher H3 acetylation in MCF-7 compared to MCF-10A cells (Sun et al., 2014). We utilized publicly available gene expression data to investigate why these cell lines may respond differently to the same doses of TSA. Consistently across two independent studies [GSE59734 (Horton et al., 2015) and GSE96860 (Xi et al., 2018)], MCF-7 cells expressed higher levels of histone acetyltransferases (HATs) and lower levels of HDACs than MCF-10A cells (**Fig. 1g**). This initial expression state difference between these cell lines is consistent with MCF-7 cells responding more strongly to HDACi treatment as the smaller pool of HDACs is easily inhibited and the abundant HATs shift the balance toward higher acetylation. MCF-7 cells have also been shown to have

### Transient TSA treatment increases X-ray-induced DNA damage signals across multiple cell types

We next tested whether the increased acetylation and chromatin decompaction caused by the 2-hour 500 nM TSA treatment would enhance cancer and non-cancer cell radiosensitivity to ionizing radiation (IR). We first used immunofluorescence to label the DNA damage marker γ-H2AX (phosphorylation of the histone variant H2AX) as a proxy for the amount of DNA damage present in each condition. DNA damage signals were not induced significantly by 500 nM TSA treatment for 2 h in either MCF-10A or MCF-7 cells (**Fig. 2a and b, Supplementary Fig. S4**). To determine the influence of TSA on the amount of radiation-induced DNA damage, we next examined the γH2AX foci 30 minutes after 5 Gy X-ray exposure in both MCF-10A and MCF-7 cells pretreated with DMSO or 500 nM TSA for 2 h. In both cancer and healthy cell lines, this high dose of X-ray irradiation produced a significantly higher level of γH2AX signal when cells were pretreated with TSA, as compared to X-ray treatment alone **(Fig. 2a and b)**. This result was corroborated in another pair of cancer and non-cancer cell lines: A375 melanoma cells and BJ5-ta skin fibroblasts both show enhanced γH2AX signal after 5 Gy X-ray when cells are pre-treated with 500 nM TSA for two hours (**Fig. 2c and d).**

**Figure 2.**
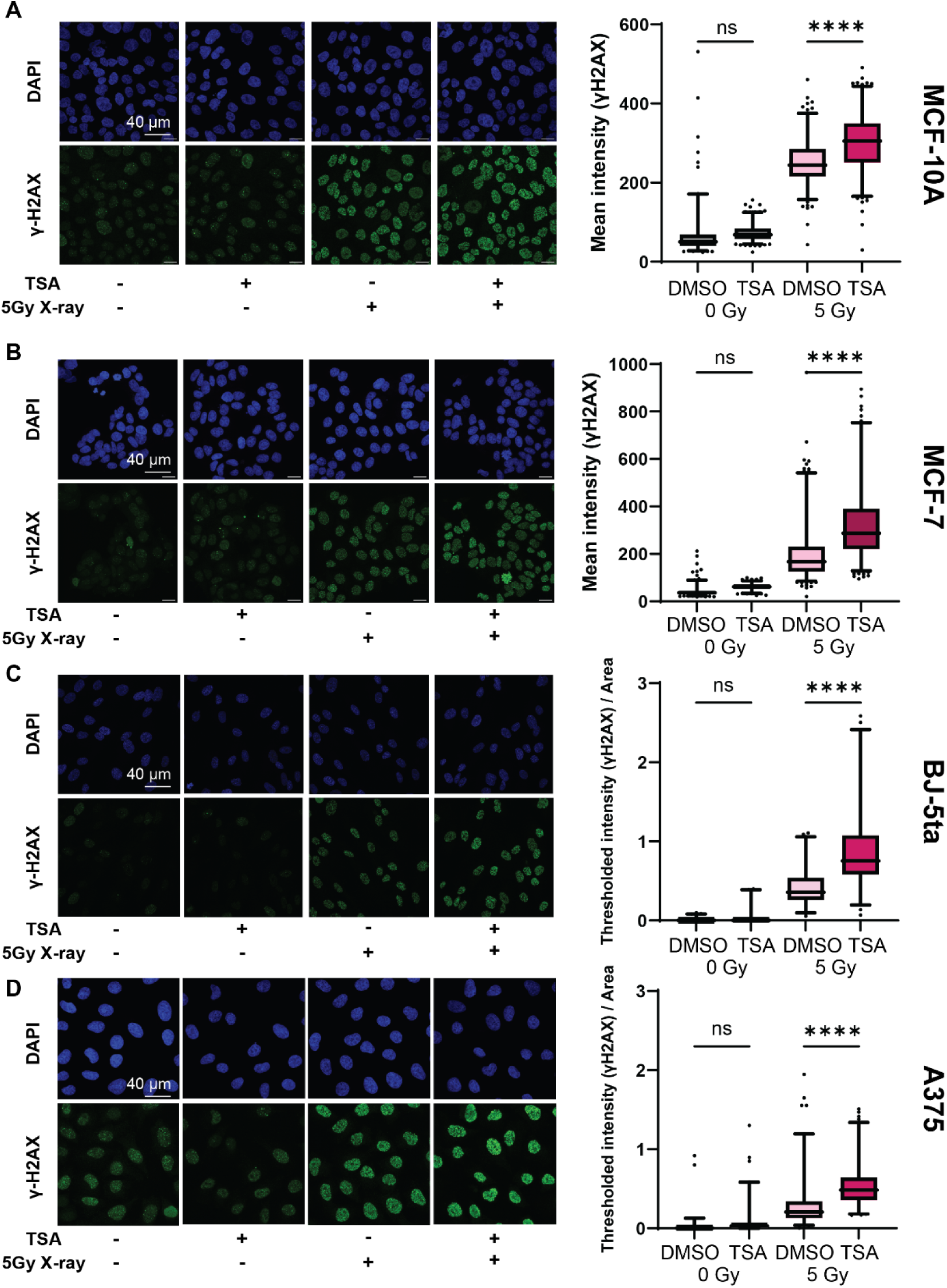
Transient TSA treatment increases DNA damage signals induced by 5 Gy X-ray across both cancer and non-cancerous cell types. **(A)** MCF-10A cells were pretreated with 500 nM TSA for 2 h, then irradiated with 5 Gy X-rays then stained with DAPI (Blue) and anti-γH2AX (Green) 30 minutes after radiation. (Left) Representative images are shown for images of cells with and without TSA treatment and with and without X-ray. Scale bar: 40 µm (shown in top left image, same for all other panels). (Right) Quantification of average γH2AX intensity per nucleus. 5 fields imaged per condition, N=280 – 376 nuclei per condition. For all boxplots, boxes show 25^th^ percentile, median, and 75^th^ percentile while whiskers show 2.5-97.5% range with outliers as points. **** p < 0.0001, ns: p>0.01, one-way ANOVA with multiple comparisons. **(B)** Same as (a) for MCF-7 cells. (N = 282-437 nuclei per condition) **(C)** Same as (a) for BJ-5ta cells. Quantification was calculated on thresholded images to eliminate background signal and then divided by nucleus area. (N=93-120 nuclei per condition) **(D)** Same as (a) for A375 cells. Quantification was calculated on thresholded images to eliminate background signal and then divided by nucleus area. (N = 156-204 nuclei per condition).

Previous literature has shown that 24 hour pre-treatment with HDACi in these same A375 melanoma cells leads to downregulation of key DNA repair genes such as Ku70, Ku86, and DNA-PKcs, which are all involved in the non homologous end joining pathway (Munshi et al., 2005, Munshi et al., 2006). We examined publicly available transcriptomic data to evaluate whether brief TSA treatment would also cause downregulation of these DNA repair genes. Two hour treatment with 500 nM TSA in A375 cells (GSE275755, (Vinayak et al., 2025)) does not significantly change the transcription of Ku70, Ku86, or DNA-PKcs (**Supplementary Fig. S5a**). In fact, no DNA damage or DNA repair pathways are found among functional annotations enriched for genes up-or down-regulated by 2 hr TSA treatment in A375 cells (**Supplementary Fig. S5b**). Existing RNA-seq data for 6 hour TSA treatment in MCF7 cells (GSE252117, (Rogers et al., 2024)) also shows no change in Ku70, Ku86, or DNA-PKcs (**Supplementary Fig. S5c**). Genes annotated as involved in cellular response to DNA damage stimulus are found among the set downregulated by 6 h TSA, but a closer look at these 32 genes reveals that they are involved in gene expression regulation and chromatin organization rather than core DNA repair pathways (**Supplementary Fig. S5d,e**). Relatedly, previous literature shows that while 24 or 48 hour treatment with HDACi caused reduced ATM expression in MCF-7 cells, 1 hour treatment had no effect (Thurn et al., 2013). Overall, these results suggest that treatment with TSA for a few hours has minimal effect on the expression of DNA repair genes, so the increased γH2AX signal after TSA treatment and 5 Gy X-ray exposure in all these cell types is more likely due to the initial effects of chromatin decompaction and acetylation rather than gene expression changes.

### Sensitizing effect of brief TSA treatment is cell-type specific at lower radiation doses

While the effect of TSA on the response to 5 Gy X-ray was consistent across cell types, we observed different responses at a lower 1 Gy X-ray dose **(Figure 3)**. In non-cancerous MCF-10A cells, pretreatment with TSA did not increase vulnerability to DNA damage after 1 Gy irradiation. In contrast, DNA damage signals produced by 1 Gy irradiation increased significantly when MCF-7 breast cancer cells were pretreated with TSA (**Fig. 3a-c, Supplementary Fig. S4d-e**). In A375 cells, increased DNA damage after 2 hours of TSA treatment was only observed at 5 Gy, not 1 Gy or 2 Gy doses. This is in contrast with published data from 24 h HDACi treatment of A375 cells, which show enhanced γH2AX signals with 2 Gy treatment (Munshi et al., 2005). This suggests that there are cell type specific properties that influence the effect of brief TSA treatment on DNA damage response at lower radiation doses.

**Figure 3.**
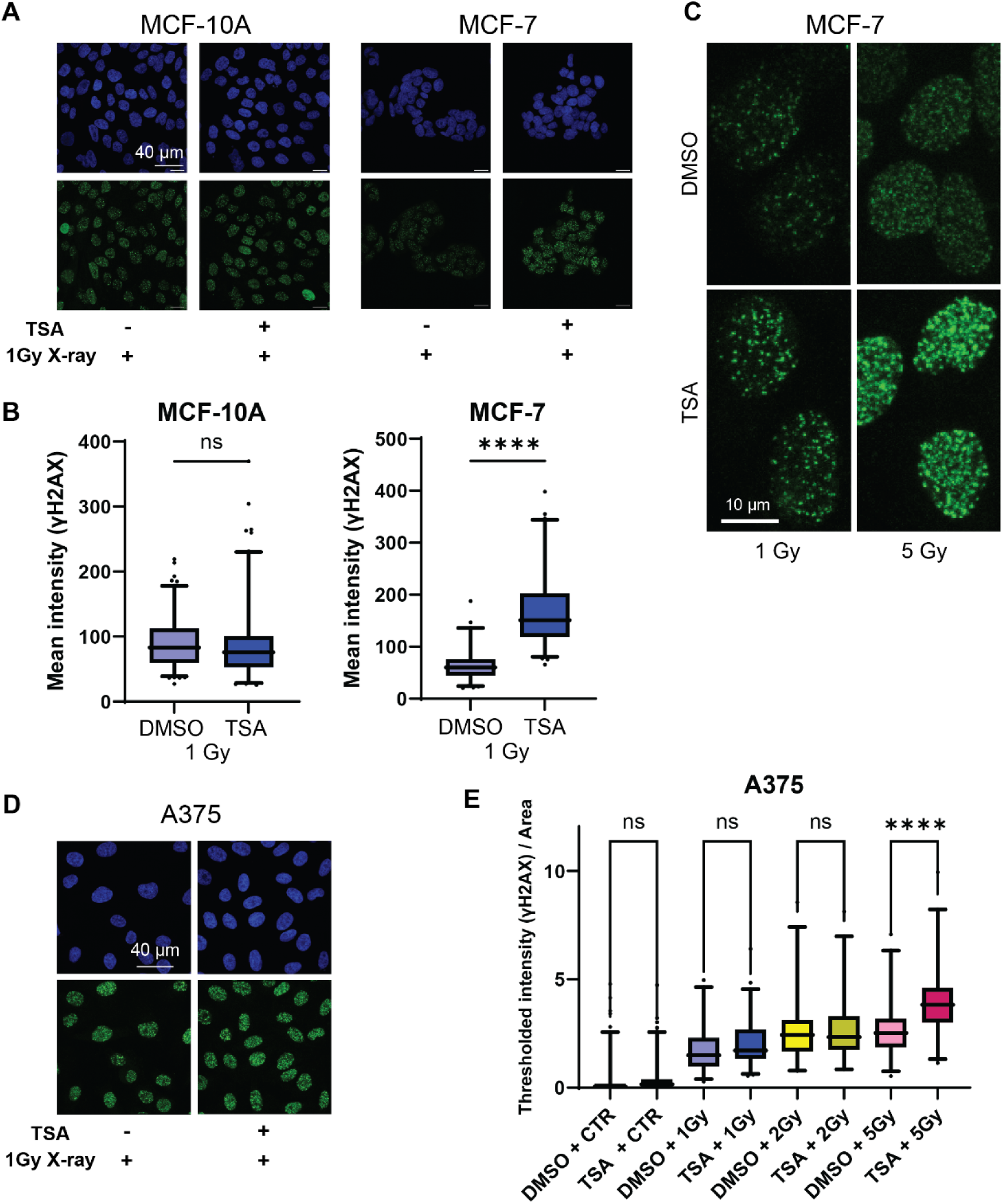
Effects of TSA on damage induced by lower radiation doses is cell type dependent. **(A)** MCF-10A (left) and MCF-7 (right) cells were pretreated with 500 nM TSA for 2 h, then irradiated with 1 Gy X-rays then stained with DAPI (Blue) and anti-γH2AX (Green) 30 minutes after radiation. Representative images are shown for images of cells with and without TSA treatment and with and without 1 Gy X-ray. Scale bar: 40 µm (shown in top left image, same for all other panels). **(B)** Quantification of average γH2AX intensity per nucleus. 5 fields imaged per condition, N=159 – 249 nuclei per condition. Boxes show 25^th^ percentile, median, and 75^th^ percentile while whiskers show 2.5-97.5% range with outliers as points. **** p < 0.0001, ns: p>0.01, one-way ANOVA with multiple comparisons. **(C)** Higher resolution, zoomed-in images of example nuclei showing enhanced DNA damage signaling after TSA treatment for both 1 and 5 Gy X-ray exposure in MCF-7 cells as compared to DMSO treatment. **(D)** Same as (A) but for 1 Gy X-ray treatment and 2 hr TSA treatment of A375 cells. **(E)** Quantification of the sum of thresholded γH2AX intensity per nucleus with or without 2 hr TSA treatment across a range of X-ray dose. (N=71-229 nuclei per condition) **** p < 0.0001, one-way ANOVA with multiple comparisons.

### Transient TSA treatment leads to altered long term radiation survival in both healthy and cancer cells

To compare the longer term response to combined TSA and X-ray treatment, we first measured the decline in γH2AX intensity over 24 h of repair time. In both MCF-7 and MCF-10a cells, despite increased damage signals after TSA treatment followed by X-ray at the 30 minute timepoint, TSA treated cells had repaired their DNA damage as much as DMSO treated cells by 24 hours. This suggests that there is not a decrease in repair rate due to 2 hour TSA treatment, but instead only an increase in initial DNA damage. Previous literature has sometimes stated that longer term HDACi treatment slows DNA repair because more damage remains at 24 hours after irradiation in HDACi treated cells. However, reanalyzing published data (Munshi et al., 2005) in A375 cells reveals that the repair rate (rate of decline in γH2AX over time) is not slower in HDACi treated cells **(Supplementary Figure S6a)**, again emphasizing the importance of the amount of initial damage in the longer term effects.

We next used a clonogenic assay to determine how the increased initial DNA damage observed after TSA treatment and irradiation would the cells’ long-term proliferative capacity. MCF-10A and MCF-7 cells were pretreated with TSA for 2 hours, then the TSA was washed out prior to 0, 1, 2 and 5 Gy X-ray irradiation followed by colony growth for 14 days. Raw measurements of total colony area in both replicates showed that 2 hour TSA treatment alone had no effect on colony growth **(Supplementary Fig. S6b).** Clonogenic survival was reduced in both MCF-10A and MCF-7 cells exposed to increasing doses of radiation after TSA treatment (**Figure 4b-c)**. The sensitizer enhancement ratios (SER) calculated from linear-quadratic fits were on the lower end of the range (1.24-3.21) described after extended (18 h) TSA treatment across numerous cell lines. Since MCF-10A cells grew in a spreading pattern, using a percent colony area quantification approach (Guzmán et al., 2014) more clearly revealed the radiosensitizing effect of TSA (SER = 1.72) on those cells **(Supplementary Fig. S6c)**, while this approach was not as clear for MCF-7 cells, where effects were more clearly seen by counting colonies with 50+ cells.

**Figure 4.**
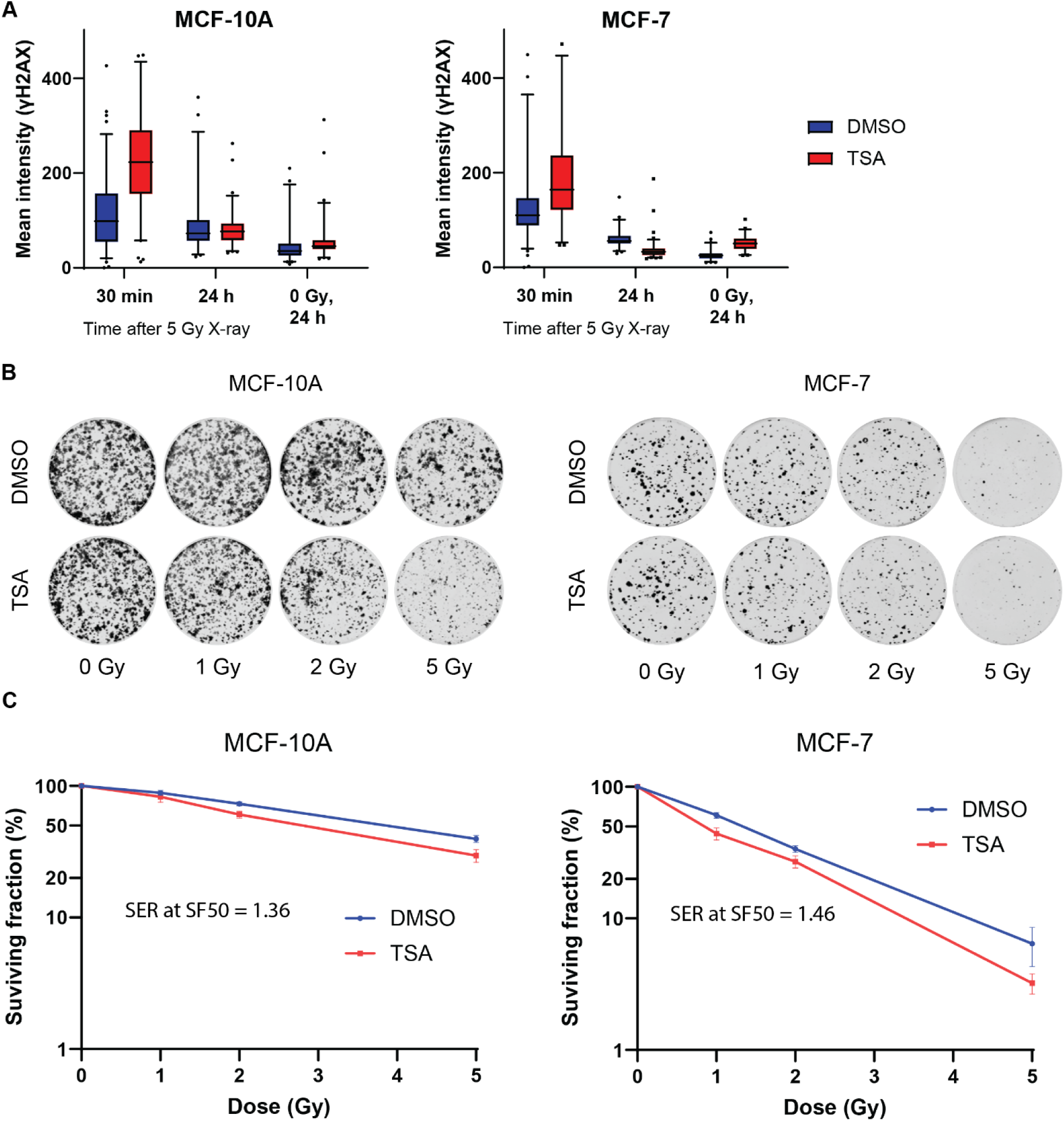
Transient TSA pre-treatment before radiation exposure has only modest effects on long term cellular phenotypes. **(A)** MCF-10A (left) and MCF-7 (right) cells were pretreated with or without 500 nM TSA for 2 h followed by 5 Gy X-ray irradiation. Boxplots show quantified mean γH2Ax per nucleus 30 min and 24 h after X-ray. The 24 h timepoint after no irradiation is shown as a no damage comparison. Boxes show 25^th^ percentile, median, and 75^th^ percentile while whiskers show 2.5-97.5% range with outliers as points. **(B)** MCF-10A (left) and MCF-7 (right) cells were pretreated with or without 500 nM TSA for 2 h, followed by 0, 1, 2, 5 Gy radiation. Representative images of colony formation are shown 14 days after indicated treatments. **(C)** The number of colonies with 50+ cells in each treatment from (B) were quantified and then normalized to the number of colonies in samples without radiation. *n = 2* biological replicates (with three technical replicates per biological replicate). Bars show mean +/- SEM averaged across all replicates. The sensitizer enhancement ratio (SER) at survival fraction 50% was calculated from a linear-quadratic fit. Data were also quantified as % area covered, as shown in Supplementary Fig S6.

## Discussion

Toward the goal of preferentially sensitizing cancer cells to radiation therapy, it is important to understand the mechanisms by which radiosensitizing agents work and whether such agents act specifically on cancer cells or certain cell types. In this study, we used TSA, a well-known HDAC inhibitor which preserves the acetylation on histone tails and increases chromatin accessibility, to increase the radiosensitivity of cells in culture. HDAC inhibitors in general, and TSA in particular, have been extensively studied as tumor inhibitors and enhancers of radiosensitivity (Ma et al., 2015, You and Park, 2013, Kim et al., 2004, Kim et al., 2005, Flatmark et al., 2006, Jia et al., 2017, Wang et al., 2021). However, the vast majority of studies only consider effects of TSA after long term (16-48 h) treatment, making it difficult to distinguish the specific effect of increased histone acetylation and chromatin accessibility from the many other effects of TSA, including its documented effects on the cell cycle (Kim et al., 2005, Zhang et al., 2009, Sun et al., 2014, Jia et al., 2017, Wang et al., 2021). In contrast, we found that a 2-hour treatment with TSA produced substantial increases in histone acetylation without affecting cell viability or proliferation. Further, many studies do not evaluate a matched non-cancerous cell control, leaving it unclear whether the radiosensitizing effects of TSA are cancer-specific or generally true for all human cells. Therefore, in this study, we focused on a transient TSA pre-treatment (2 h) and compared effects on both cancer and non-cancerous cell lines.

Our data show that 2 hour TSA pretreatment increases initial DNA damage signals after 5 Gy X-ray irradiation not only in breast cancer and melanoma cells but also in non-tumorigenic breast epithelial cells and skin fibroblasts. This response is therefore more likely to be a ubiquitous physical effect of opening and acetylating chromatin rather than a cell type specific effect based on the particular initial chromatin state or repair factor expression levels. This result is important both because it reveals that increased histone acetylation influences X-ray induced DNA damage formation generally across cell types and because it cautions against studying factors that radiosensitize cancer cells without considering potentially similar effects on healthy cells.

Many previous studies on the effects of HDACi treatment on radiosensitivity focus on the downstream changes in expression of DNA damage repair proteins caused by long term HDACi treatment (Zhang et al., 2007, Munshi et al., 2005). However, we find that these genes do not change expression after short term TSA treatment and therefore cannot explain the increase in initial DNA damage signals that we observe. Therefore, this increase in DNA damage signal right after irradiation is more likely to stem from HDACi impacts on the chromatin state itself. Some part of this effect may arise from the role of histone acetylation in DNA damage repair signaling. Extensive literature over the past 4 decades has revealed a complex spatiotemporal acetylation response to DNA damage that involves both acetylation signals to recruit or prevent certain repair enzymes, to relax the chromatin for repair, and to influence repair pathway choice (Aricthota et al., 2022, Ramanathan and Smerdon, 1986, Zhang et al., 2021, Seane et al., 2024b). The acetylation of certain histone sites, such as H3K14Ac, are beneficial to cell survival and repair because they promote spreading of ATM activation and γH2AX. In contrast, high levels of H3K9Ac inhibit ATM activation and increase radiosensitivity (Meyer et al., 2016, Tjeertes et al., 2009). However, it is important to note that our experiments remove the TSA treatment before irradiation, so we are influencing the initial state of the chromatin more than downstream acetylation responses that occur as part of the DNA repair process. Previous results show that applying HDACi only after irradiation does not have as much effect as pre-treatment (Kim et al., 2013). Therefore, our results likely reflect mechanisms by which initially more open chromatin results in more DNA damage. More open chromatin can be more vulnerable to radiation damage than compact chromatin because DNA-bound proteins and nucleosomes protect DNA from damage by OH radicals and compacted chromatin has less surrounding water that can be converted into radicals (Warters and Lyons, 1992, Nygren et al., 1995).

While increased DNA damage after brief TSA treatment was a cell type invariant feature at a 5 Gy dose, our study further showed that the effect of TSA is cell type specific at a lower radiation dose (1 Gy), increasing damage signals in MCF-7, but not MCF-10A or A375 cells. We posit that higher radiation doses saturate the potential damage sites and response machinery, resulting in a more similar effect across cell types. But, the effects of lower radiation doses are more influenced by cell type specific differences. Our analysis of previously published data on differences in the initial state of these cell types suggests that this cell type specificity could stem from higher initial levels of HATs and lower levels of HDACs in MCF7 cells than MCF10a and a concomitant stronger acetylation response to TSA in MCF7 cells. Indeed, altered expression of HDACs has previously been linked to differential HDACi and DNA damage responses in breast cancer (Shan et al., 2017). Our results suggest that the relative HAT/HDAC levels in different cell types will influence their DNA damage response in the presence of HDACi, particularly at lower doses of radiation.

Beyond initial acetylation state, differences in how each cell type responds to radiation could be another potential contributing factor to differential radiosensitization by TSA to the same dose of X-ray. Publicly available transcriptomic data for MCF-7 and MCF-10A cells measured 24 hours after 5 Gy [GSE59734, (Horton et al., 2015)] or 9 Gy [GSE108962,(Bravatà et al., 2018)] radiation shows cell-type specific differences in radiation-responsive and repair pathway gene expression **(Supplementary Fig. S7)**. We found that MCF-7 cells more strongly upregulate mismatch repair, homologous recombination, and base excision repair genes in response to X-ray, which could account for their more sensitive γH2AX response to lower doses of X-ray combined with TSA and differences in their long term survival.

How initial amounts of DNA damage translate into cell survival depends on numerous DNA repair and downstream signaling factors. Overall, 2 hour TSA treatment prior to X-ray irradiation in MCF-7 and MCF-10A cells produced a long term radiosensitization effect that was on the lower end of the effects previously seen across cell lines with an extended (18 h) TSA treatment (Kim et al., 2010). However, the SER values are comparable with effects seen by 24 h treatment with the HDACi CUDC-101 in MCF-7 and MCF-10A cells (Seane et al., 2024a). It is noteworthy that this radiosensitizing effect is produced without the TSA treatment on its own causing cell viability effects, unlike longer TSA treatments. Our observations suggest that the effects of longer term TSA treatment on DNA repair protein expression and the cell cycle are not essential to explain the HDACi-mediated increase in DNA damage, and this increase in damage can on its own affect cell viability, though altering repair and cell cycle with extended HDACi treatment can enhance cell killing long term. Importantly, our results show that non-cancerous MCF10a cells have notably reduced long term survival in combined TSA and X-ray treatment, perhaps even more so than the cancerous cells. These results show that initial DNA damage amounts do not simply convert into long term survival rates (Yasushi et al., 2007) and that comparing effects on non-cancerous cells is important when evaluating a radiosensitizing treatment.

Overall, our results suggest that initial histone acetylation and chromatin decompaction play an important part in the mechanism of TSA radiosensitization and generally increase DNA damage across cell types under high doses of radiation. Our study suggests that responses to lower doses of radiation, where we saw more cell type specific divergence should be further explored. Indeed, most studies reporting radiosensitizing effects of TSA see the most obvious effects in the 4-8 Gy dose range (Kim et al., 2010). But, given that many therapeutic radiation doses are applied in the 1-2 Gy dose range, it is important to understand differential responses to radiosensitizers at these lower doses. Given that just two hours of TSA treatment can elicit radiosensitizing responses suggests it is also important to consider whether treatment timing or local application of HDACis could maximize radiosensitization while minimizing effects of the HDACi treatment alone.

## AUTHOR CONTRIBUTIONS

H.L., S.D., and R.P.M. conceived the study. H.L., C.M., and L.Y. performed experiments and data analyses and prepared figures. R.P.M. supervised and guided the work. S.D. and D.P. advised on clonogenic assays and data interpretation. H.L. and R.P.M. wrote the manuscript with input from all authors.

## ACKNOWLEDGEMENTS

The authors would like to thank all the members of McCord lab for their fruitful suggestions in developing the research project. The authors would like to acknowledge the University of Tennessee Advanced Microscopy and Imaging Center (RRID:SCR_028273) for instrument use and scientific and technical assistance.

## SUPPLEMENTARY MATERIAL

The supplementary material associated with the paper can be found and accessed online.

## References

Alao, J. P., Lam, E. W. F., Ali, S., Buluwela, L., Bordogna, W., Lockey, P., Varshochi, R., Stavropoulou, A. V., Coombes, R. C. & Vigushin, D. M. 2004. Histone Deacetylase Inhibitor Trichostatin A Represses Estrogen Receptor α-Dependent Transcription and Promotes Proteasomal Degradation of Cyclin D1 in Human Breast Carcinoma Cell Lines. Clinical Cancer Research, 10, 8094–8104.

Aricthota, S., Rana, P. P. & Haldar, D. 2022. Histone acetylation dynamics in repair of DNA double-strand breaks. Frontiers in Genetics, Volume 13–2022.

Bouyahya, A., El Omari, N., Bakha, M., Aanniz, T., El Menyiy, N., EL Hachlafi, N., El Baaboua, A., El-Shazly, M., Alshahrani, M. M., Al Awadh, A. A., Lee, L.-H., Benali, T. & Mubarak, M. S. 2022. Pharmacological Properties of Trichostatin A, Focusing on the Anticancer Potential: A Comprehensive Review. Pharmaceuticals [Online], 15.

Brambilla, F., Garcia-Manteiga, J. M., Monteleone, E., Hoelzen, L., Zocchi, A., Agresti, A. & Bianchi, M. E. 2020. Nucleosomes effectively shield DNA from radiation damage in living cells. Nucleic Acids Research, 48, 8993–9006.

Bravatà, V., Cava, C., Minafra, L., Cammarata, F. P., Russo, G., Gilardi, M. C., Castiglioni, I. & Forte, G. I. 2018. Radiation-Induced Gene Expression Changes in High and Low Grade Breast Cancer Cell Types. International Journal of Molecular Sciences [Online], 19.

Brix, N., Samaga, D., Hennel, R., Gehr, K., Zitzelsberger, H. & Lauber, K. 2020. The clonogenic assay: robustness of plating efficiency-based analysis is strongly compromised by cellular cooperation. Radiation Oncology, 15, 248.

Cao, R., Wang, L., Wang, H., Xia, L., Erdjument-Bromage, H., Tempst, P., Jones, R. S. & Zhang, Y. 2002. Role of histone H3 lysine 27 methylation in Polycomb-group silencing. Science, 298, 1039–43.

Citrin, D. E. & Mitchell, J. B. 2014. Altering the response to radiation: sensitizers and protectors. Semin Oncol, 41, 848–59.

Cowell, I. G., Sunter, N. J., Singh, P. B., Austin, C. A., Durkacz, B. W. & Tilby, M. J. 2007. γH2AX Foci Form Preferentially in Euchromatin after Ionising-Radiation. PLOS One, 2, e1057.

Das, P., Golloshi, R., Mccord, R. P. & Shen, T. 2020. Using contact statistics to characterize structure transformation of biopolymer ensembles. Phys Rev E, 101, 012419.

De, K., Grubb, T. M., Zalenski, A. A., Pfaff, K. E., Pal, D., Majumder, S., Summers, M. K. & Venere, M. 2019. Hyperphosphorylation of CDH1 in Glioblastoma Cancer Stem Cells Attenuates APC/C(CDH1) Activity and Pharmacologic Inhibition of APC/C(CDH1/CDC20) Compromises Viability. Mol Cancer Res, 17, 1519–1530.

Falk, M., Lukášová, E. & Kozubek, S. 2008. Chromatin structure influences the sensitivity of DNA to γ-radiation. Biochimica et Biophysica Acta (BBA) - Molecular Cell Research, 1783, 2398–2414.

Ferrari, K. J., Scelfo, A., Jammula, S., Cuomo, A., Barozzi, I., Stutzer, A., Fischle, W., Bonaldi, T. & Pasini, D. 2014. Polycomb-dependent H3K27me1 and H3K27me2 regulate active transcription and enhancer fidelity. Mol Cell, 53, 49–62.

Flatmark, K., Nome, R. V., Folkvord, S., Bratland, A., Rasmussen, H., Ellefsen, M. S., Fodstad, O. & Ree, A. H. 2006. Radiosensitization of colorectal carcinoma cell lines by histone deacetylase inhibition. Radiat Oncol, 1, 25.

Franken, N. A., Rodermond, H. M., Stap, J., Haveman, J. & Van Bree, C. 2006. Clonogenic assay of cells in vitro. Nat Protoc, 1, 2315–9.

Gingrich, P. W., Chitsazi, R., Biswas, A., Jiang, C., Zhao, L., Tym, J. E., Brammer, K. M., Li, J., Shu, Z., Maxwell, D. S., Tacy, J. A., Mica, I. L., Darkoh, M., Di Micco, P., Russell, K. P., Workman, P. & Al-Lazikani, B. 2025. canSAR 2024-an update to the public drug discovery knowledgebase. Nucleic Acids Res, 53, D1287–D1294.

Guzmán, C., Bagga, M., Kaur, A., Westermarck, J. & Abankwa, D. 2014. ColonyArea: An ImageJ Plugin to Automatically Quantify Colony Formation in Clonogenic Assays. PLOS One, 9, e92444.

He, G., Wang, Y., Pang, X. & Zhang, B. 2014. Inhibition of autophagy induced by TSA sensitizes colon cancer cell to radiation. Tumour Biol, 35, 1003–11.

Heo, S. J., Song, K. H., Thakur, S., Miller, L. M., Cao, X., Peredo, A. P., Seiber, B. N., Qu, F., Driscoll, T. P., Shenoy, V. B., Lakadamyali, M., Burdick, J. A. & Mauck, R. L. 2020. Nuclear softening expedites interstitial cell migration in fibrous networks and dense connective tissues. Sci Adv, 6, eaax5083.

Horton, J. K., Siamakpour-Reihani, S., Lee, C. T., Zhou, Y., Chen, W., Geradts, J., Fels, D. R., Hoang, P., Ashcraft, K. A., Groth, J., Kung, H. N., Dewhirst, M. W. & Chi, J. T. 2015. FAS Death Receptor: A Breast Cancer Subtype-Specific Radiation Response Biomarker and Potential Therapeutic Target. Radiat Res, 184, 456–69.

Jia, L., Zhang, S., Huang, Y., Zheng, Y. & Gan, Y. 2017. Trichostatin A increases radiosensitization of tongue squamous cell carcinoma via mir-375. Oncol Rep, 37, 305–312.

Karagiannis, T. C., N., H. K. & and El-Osta, A. 2005. The histone deacetylase inhibitor, trichostatin A, enhances radiation sensitivity and accumulation of gammaH2A.X. Cancer Biology & Therapy, 4, 787–793.

Kim, I. A., Kim, I. H., Kim, H. J., Chie, E. K. & Kim, J. S. 2010. HDAC inhibitor-mediated radiosensitization in human carcinoma cells: a general phenomenon? J Radiat Res, 51, 257–63.

Kim, I. A., Kim, J. H., Shin, J. H., Kim, I. H., Kim, J. S., Wu, H. G., Chie, E. K., Kim, Y. H., Kim, B. K., Hong, S., Park, S. W., Ha, S. W. & Park, C. I. 2005. A histone deacetylase inhibitor, trichostatin A, enhances radiosensitivity by abrogating G2/M arrest in human carcinoma cells. Cancer Res Treat, 37, 122–8.

Kim, J. H., Kim, I. H., Shin, J. H., Kim, H. J. & Kim, I. A. 2013. Sequence-Dependent Radiosensitization of Histone Deacetylase Inhibitors Trichostatin A and Sk-7041. Cancer Res Treat, 45, 334–342.

Kim, J. H., Shin, J. H. & Kim, I. H. 2004. Susceptibility and radiosensitization of human glioblastoma cells to trichostatin A, a histone deacetylase inhibitor. Int J Radiat Oncol Biol Phys, 59, 1174–80.

Kim, T. H., Nosella, M. L., Bolik-Coulon, N., Harkness, R. W., Huang, S. K. & Kay, L. E. 2023. Correlating histone acetylation with nucleosome core particle dynamics and function. Proc Natl Acad Sci U S A, 120, e2301063120.

Kong, W. Y., Yee, Z. Y., Mai, C. W., Fang, C.-M., Abdullah, S. & Ngai, S. C. 2019. Zebularine and trichostatin A sensitized human breast adenocarcinoma cells towards tumor necrosis factor-related apoptosis inducing ligand (TRAIL)-induced apoptosis. Heliyon, 5, e02468.

Kurashige, T., Shimamura, M. & Nagayama, Y. 2016. Differences in quantification of DNA double-strand breaks assessed by 53BP1/γH2AX focus formation assays and the comet assay in mammalian cells treated with irradiation and N-acetyl-L-cysteine. Journal of Radiation Research, 57, 312–317.

Lachner, M., O’carroll, D., Rea, S., Mechtler, K. & Jenuwein, T. 2001. Methylation of histone H3 lysine 9 creates a binding site for HP1 proteins. Nature, 410, 116–20.

Ma, J., Guo, X., Zhang, S., Liu, H., Lu, J., Dong, Z., Liu, K. & Ming, L. 2015. Trichostatin A, a histone deacetylase inhibitor, suppresses proliferation and promotes apoptosis of esophageal squamous cell lines. Mol Med Rep, 11, 4525–31.

Meyer, B., Fabbrizi, M. R., Raj, S., Zobel, C. L., Hallahan, D. E. & Sharma, G. G. 2016. Histone H3 Lysine 9 Acetylation Obstructs ATM Activation and Promotes Ionizing Radiation Sensitivity in Normal Stem Cells. Stem Cell Reports, 7, 1013–1022.

Min, K. N., Cho, M. J., Kim, D.-K. & Sheen, Y. Y. 2004. Estrogen receptor enhances the antiproliferative effects of trichostatin A and Hc-toxin in human breast cancer cells. Archives of Pharmacal Research, 27, 554–561.

Munshi, A., Kurland, J. F., Nishikawa, T., Tanaka, T., Hobbs, M. L., Tucker, S. L., Ismail, S., Stevens, C. & Meyn, R. E. 2005. Histone Deacetylase Inhibitors Radiosensitize Human Melanoma Cells by Suppressing DNA Repair Activity. Clinical Cancer Research, 11, 4912–4922.

Munshi, A., Tanaka, T., Hobbs, M. L., Tucker, S. L., Richon, V. M. & Meyn, R. E. 2006. Vorinostat, a histone deacetylase inhibitor, enhances the response of human tumor cells to ionizing radiation through prolongation of γ-H2AX foci. Molecular Cancer Therapeutics, 5, 1967–1974.

Noh, H., Park, J., Shim, M. & Lee, Y. 2016. Trichostatin A enhances estrogen receptor-alpha repression in Mcf-7 breast cancer cells under hypoxia. Biochemical and Biophysical Research Communications, 470, 748–752.

Nygren, J., Ljungman, M. & Ahnström, M. 1995. Chromatin Structure and Radiation-induced DNA Strand Breaks in Human Cells: Soluble Scavengers and Dna-bound Proteins Offer a Better Protection Against Single- than Double-strand Breaks. International Journal of Radiation Biology, 68, 11–18.

Paillas, S., Then, C. K., Kilgas, S., Ruan, J.-L., Thompson, J., Elliott, A., Smart, S. & Kiltie, A. E. 2020. The Histone Deacetylase Inhibitor Romidepsin Spares Normal Tissues While Acting as an Effective Radiosensitizer in Bladder Tumors in Vivo. International Journal of Radiation Oncology*Biology*Physics, 107, 212–221.

Paldi, F., Szalay, M.-F., Dufau, S., Di Stefano, M., Reboul, H., Jost, D., Bantignies, F. & Cavalli, G. 2026. Transient histone deacetylase inhibition induces cellular memory of gene expression and 3D genome folding. Nature Genetics, 58, 404–417.

Ramanathan, B. & Smerdon, M. J. 1986. Changes in nuclear protein acetylation in u.v.-damaged human cells. Carcinogenesis, 7, 1087–1094.

Rogers, J. D., Leusch, F. D. L., Chambers, B., Daniels, K. D., Everett, L. J., Judson, R., Maruya, K., Mehinto, A. C., Neale, P. A., Paul-Friedman, K., Thomas, R., Snyder, S. A. & Harrill, J. 2024. High-Throughput Transcriptomics of Water Extracts Detects Reductions in Biological Activity with Water Treatment Processes. Environmental Science & Technology, 58, 2027–2037.

Seane, E. N., Nair, S., Vandevoorde, C., Bisio, A. & Joubert, A. 2024a. Multi-Target Inhibitor Cudc-101 Impairs DNA Damage Repair and Enhances Radiation Response in Triple-Negative Breast Cell Line. Pharmaceuticals [Online], 17.

Seane, E. N., Nair, S., Vandevoorde, C. & Joubert, A. 2024b. Mechanistic Sequence of Histone Deacetylase Inhibitors and Radiation Treatment: An Overview. Pharmaceuticals [Online], 17.

Shan, W., Jiang, Y., Yu, H., Huang, Q., Liu, L., Guo, X., Li, L., Mi, Q., Zhang, K. & Yang, Z. 2017. HDAC2 overexpression correlates with aggressive clinicopathological features and Dna-damage response pathway of breast cancer. Am J Cancer Res, 7, 1213–1226.

Stephens, A. D., Liu, P. Z., Banigan, E. J., Almassalha, L. M., Backman, V., Adam, S. A., Goldman, R. D. & Marko, J. F. 2018. Chromatin histone modifications and rigidity affect nuclear morphology independent of lamins. Mol Biol Cell, 29, 220–233.

Sun, S., Han, Y., Liu, J., Fang, Y., Tian, Y., Zhou, J., Ma, D. & Wu, P. 2014. Trichostatin A targets the mitochondrial respiratory chain, increasing mitochondrial reactive oxygen species production to trigger apoptosis in human breast cancer cells. PLoS One, 9, e91610.

Takata, H., Hanafusa, T., Mori, T., Shimura, M., Iida, Y., Ishikawa, K., Yoshikawa, K., Yoshikawa, Y. & Maeshima, K. 2013. Chromatin compaction protects genomic DNA from radiation damage. PLoS One, 8, e75622.

Tang, N., Bueno, M., Meylan, S., Incerti, S., Tran, H. N., Vaurijoux, A., Gruel, G. & Villagrasa, C. 2019. Influence of chromatin compaction on simulated early radiation-induced DNA damage using Geant4-DNA. Med Phys, 46, 1501–1511.

Thurn, K. T., Thomas, S., Raha, P., Qureshi, I. & Munster, P. N. 2013. Histone Deacetylase Regulation of Atm-Mediated DNA Damage Signaling. Molecular Cancer Therapeutics, 12, 2078–2087.

Tjeertes, J. V., Miller, K. M. & Jackson, S. P. 2009. Screen for Dna-damage-responsive histone modifications identifies H3K9Ac and H3K56Ac in human cells. The EMBO Journal, 28, 1878–1889.

Urbinati, G., Marsaud, V., Nicolas, V., Vergnaud-Gauduchon, J. & Renoir, J.-M. 2011. Liposomal trichostatin A: therapeutic potential in hormone-dependent and - independent breast cancer xenograft models. Hormone Molecular Biology and Clinical Investigation, 6, 215–225.

Vigushin, D. M., Ali, S., Pace, P. E., Mirsaidi, N., Ito, K., Adcock, I. & Coombes, R. C. 2001. Trichostatin A Is a Histone Deacetylase Inhibitor with Potent Antitumor Activity against Breast Cancer in Vivo. Clinical Cancer Research, 7, 971–976.

Vinayak, V., Basir, R., Golloshi, R., Toth, J., Sant’anna, L., Lakadamyali, M., Mccord, R. P. & Shenoy, V. B. 2025. Polymer model integrates imaging and sequencing to reveal how nanoscale heterochromatin domains influence gene expression. Nature Communications, 16, 3816.

Wang, S., Song, M. & Zhang, B. 2021. Trichostatin A enhances radiosensitivity and radiation-induced DNA damage of esophageal cancer cells. J Gastrointest Oncol, 12, 1985–1995.

Wang, Z., Zang, C., Cui, K., Schones, D. E., Barski, A., Peng, W. & Zhao, K. 2009. Genome-wide Mapping of HATs and HDACs Reveals Distinct Functions in Active and Inactive Genes. Cell, 138, 1019–1031.

Warters, R. L. & Lyons, B. W. 1992. Variation in Radiation-Induced Formation of DNA Double-Strand Breaks as a Function of Chromatin Structure. Radiation Research, 130, 309–318.

Xi, Y., Shi, J., Li, W., Tanaka, K., Allton, K. L., Richardson, D., Li, J., Franco, H. L., Nagari, A., Malladi, V. S., Coletta, L. D., Simper, M. S., Keyomarsi, K., Shen, J., Bedford, M. T., Shi, X., Barton, M. C., Kraus, W. L., Li, W. & Dent, S. Y. R. 2018. Histone modification profiling in breast cancer cell lines highlights commonalities and differences among subtypes. BMC Genomics, 19, 150.

Yan, H., Xiang, X., Chen, Q., Pan, X., Cheng, H. & Wang, F. 2018. HP1 cooperates with CAF-1 to compact heterochromatic transgene repeats in mammalian cells. Sci Rep, 8, 14141.

Yasushi, K., Jeffrey, S. M., Kenneth, L. B. & David, J. G. 2007. Relationship between Phosphorylated Histone H2AX Formation and Cell Survival in Human Microvascular Endothelial Cells (HMEC) as a Function of Ionizing Radiation Exposure in the Presence or Absence of Thiol-Containing Drugs. Radiation Research, 168, 106–114.

You, B. R. & Park, W. H. 2013. Trichostatin A induces apoptotic cell death of HeLa cells in a Bcl-2 and oxidative stress-dependent manner. Int J Oncol, 42, 359–66.

Yu, J., Mi, J., Wang, Y., Wang, A. & Tian, X. 2012. Regulation of radiosensitivity by HDAC inhibitor trichostatin A in the human cervical carcinoma cell line Hela. Eur J Gynaecol Oncol, 33, 285–90.

Zhang, F., Zhang, T., Teng, Z. H., Zhang, R., Wang, J. B. & Mei, Q. B. 2009. Sensitization to gamma-irradiation-induced cell cycle arrest and apoptosis by the histone deacetylase inhibitor trichostatin A in non-small cell lung cancer (NSCLC) cells. Cancer Biol Ther, 8, 823–31.

Zhang, J., Lu, X., Moghaddamkohi, S., Shi, L., Xu, X. & Zhu, W.-G. 2021. Histone lysine modifying enzymes and their critical roles in DNA double-strand break repair. DNA Repair, 107, 103206.

Zhang, Y., Carr, T., Dimtchev, A., Zaer, N., Dritschilo, A. & Jung, M. 2007. Attenuated DNA damage repair by trichostatin A through BRCA1 suppression. Radiat Res, 168, 115–24.

