## Supplementary Figures for "Transient histone deacetylase inhibition reveals cell type invariant and specific effects of chromatin decondensation on irradiation response"

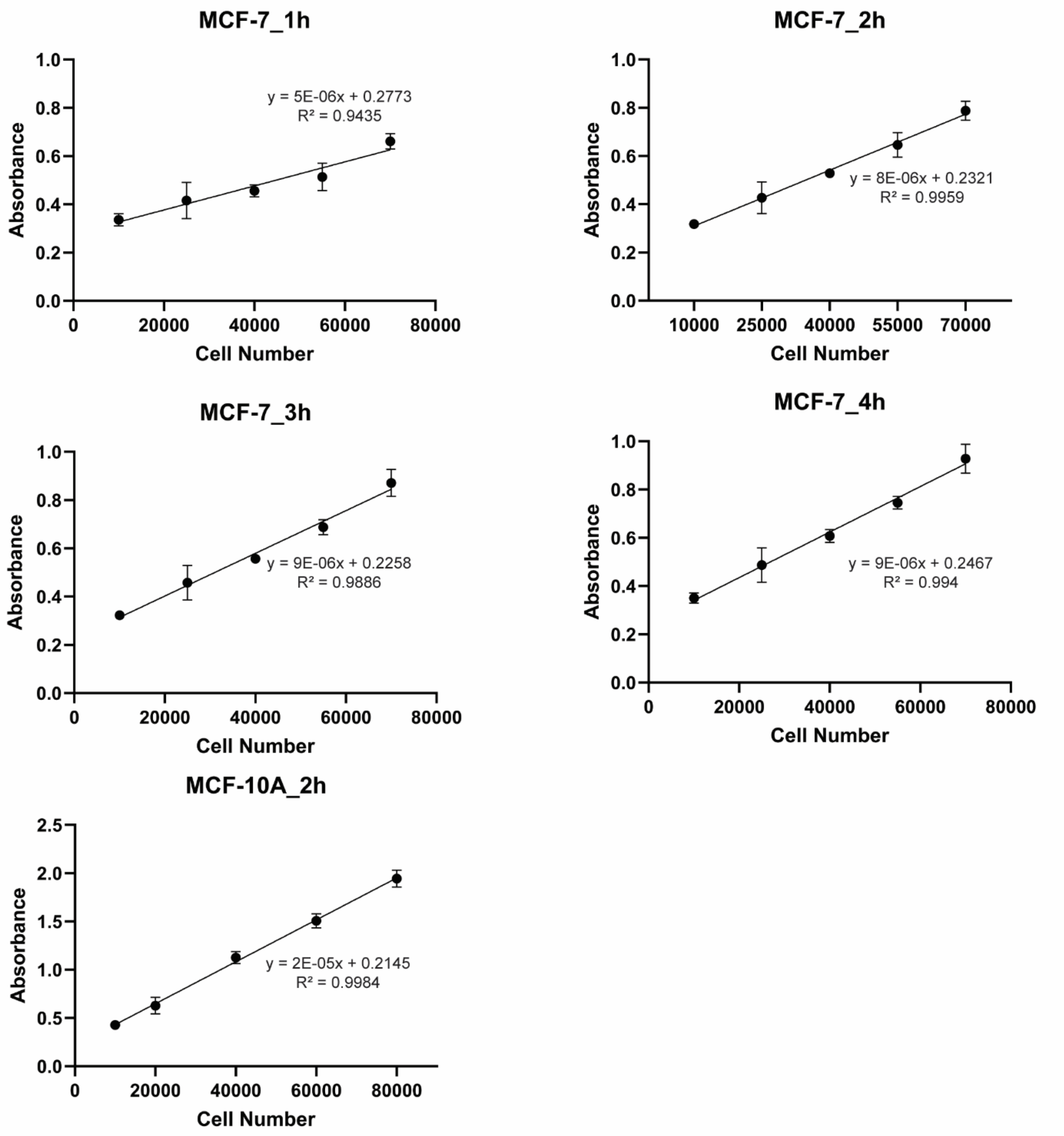


**Supplementary Figure 1. Establishing optimal MTS assay conditions.** Various numbers of MCF-10A or MCF-7 cells were seeded into wells of a 96-well plate 24 hours prior to each MTS assay. Cells were then incubated for the indicated time (1, 2, 3, or 4 h) with MTS reagents, and absorbance was measured to indicate cellular metabolic activity. Each point represents the mean ± SD of 6 replicates. The correlation coefficient indicates a linear response between the cell number and absorbance. Two-hour incubation was chosen as the optimal timing based on having the best correlation coefficient for MCF-7 and a strong correlation for MCF-10A as well.

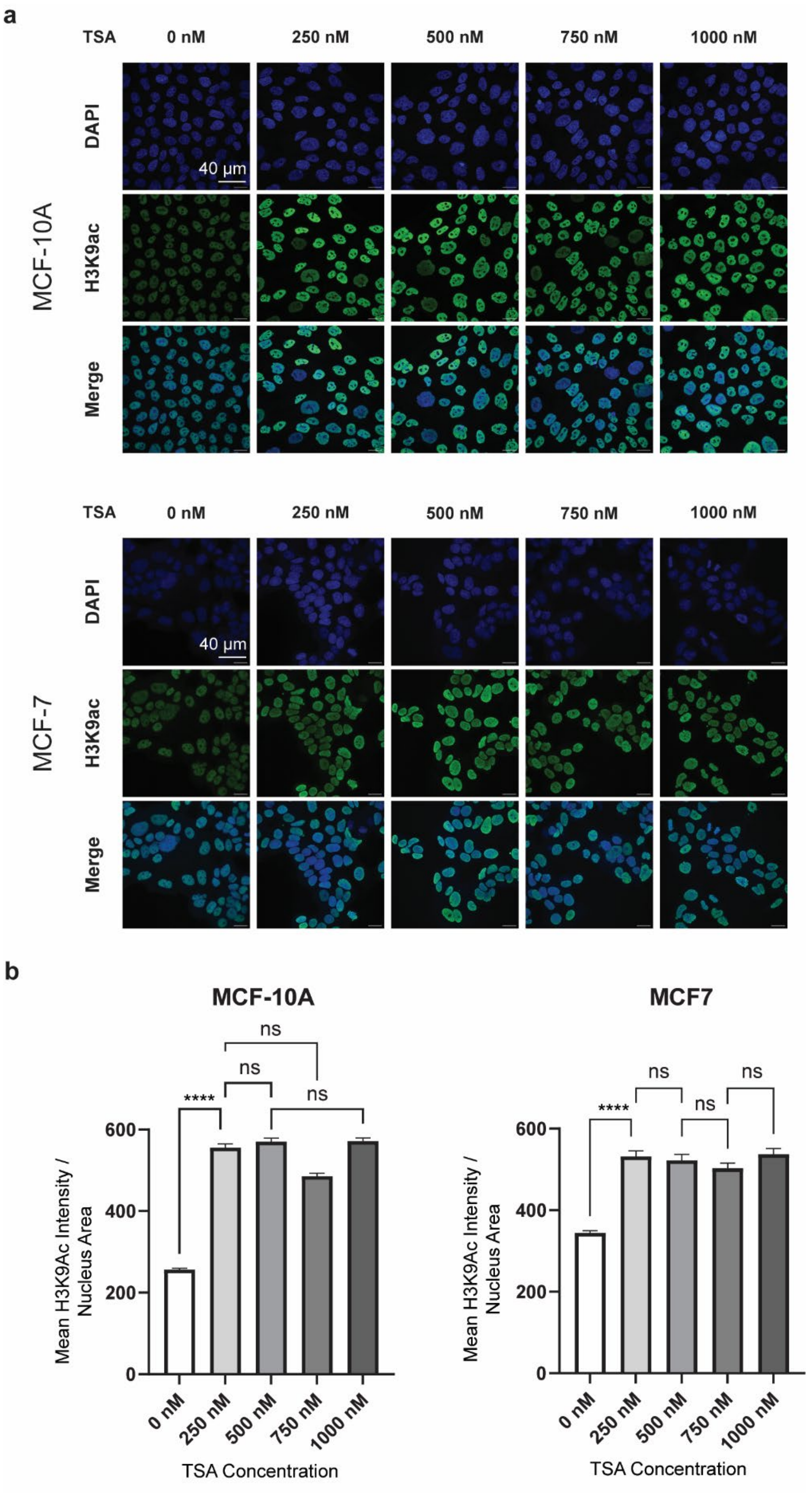


**Supplementary Figure 2. TSA enhances histone acetylation – biological replicate experiment.** This figure presents data from an independent biological replicate of the experiments shown in Figure 1c. (**a**) MCF-10A (top) and MCF-7 (bottom) cells were treated with indicated concentration of TSA (0, 250, 500, 750 and 1000 nM) for 2 h, and stained with DAPI (Blue) and anti-H3K9ac (Green) in the indicated treatments. Images are maximum projections of Z-stacks. Scale bar: 40 µm (top left image, same for all other panels). (**b**) Quantification of average H3K9ac intensity per nucleus area. Bars show mean +/- SEM averaged across all measured nuclei (5 fields imaged per condition, N=150 – 300 nuclei per condition) (**** $p < 0.0001$ for pairwise one-tailed t-tests between each increasing concentration)

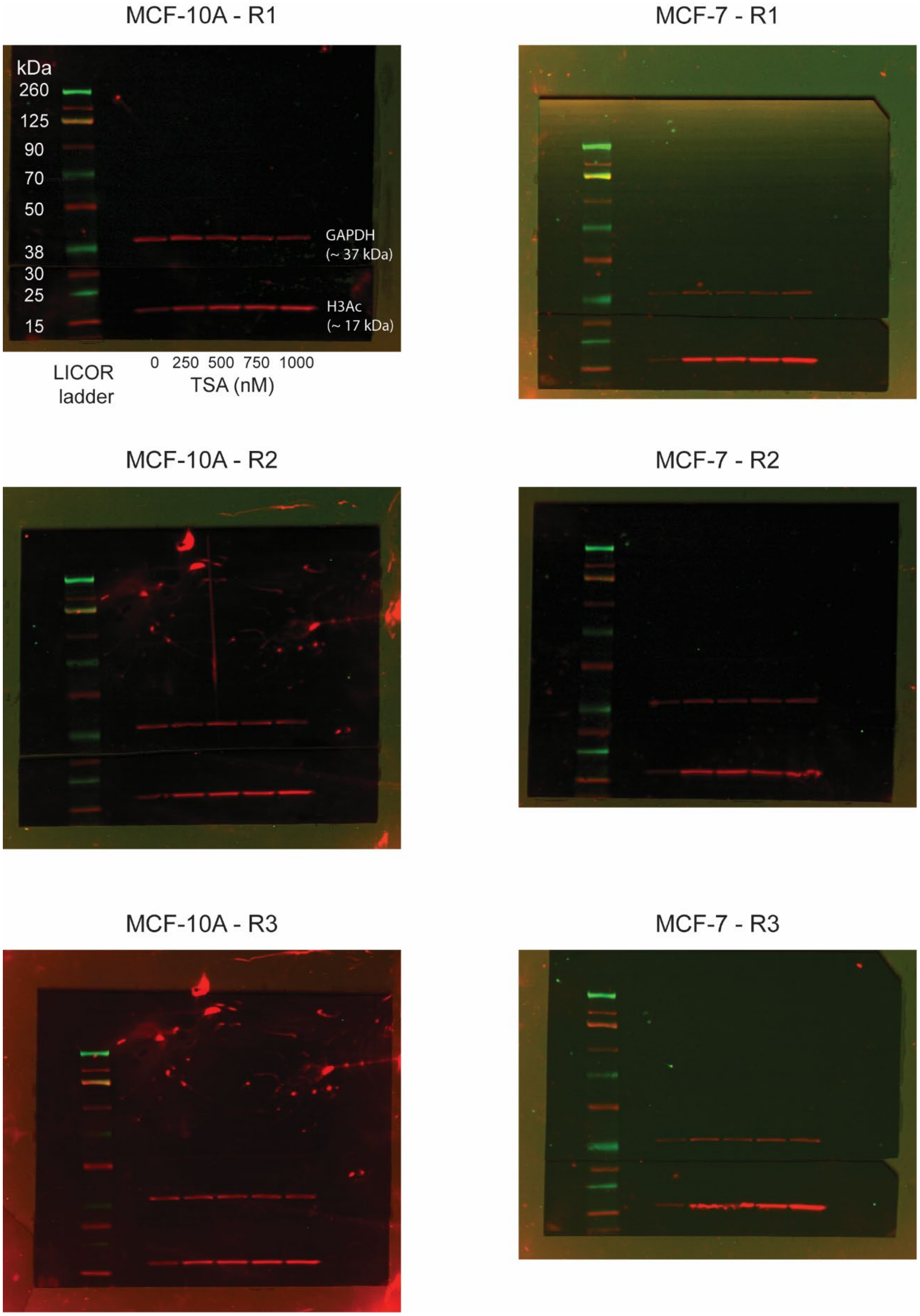


**Supplementary Figure 3. Original, uncropped blots supporting histone acetylation quantification in Figure 1e.** Three biological replicates for each cell line are shown. Labels on the top left blot apply to all images.

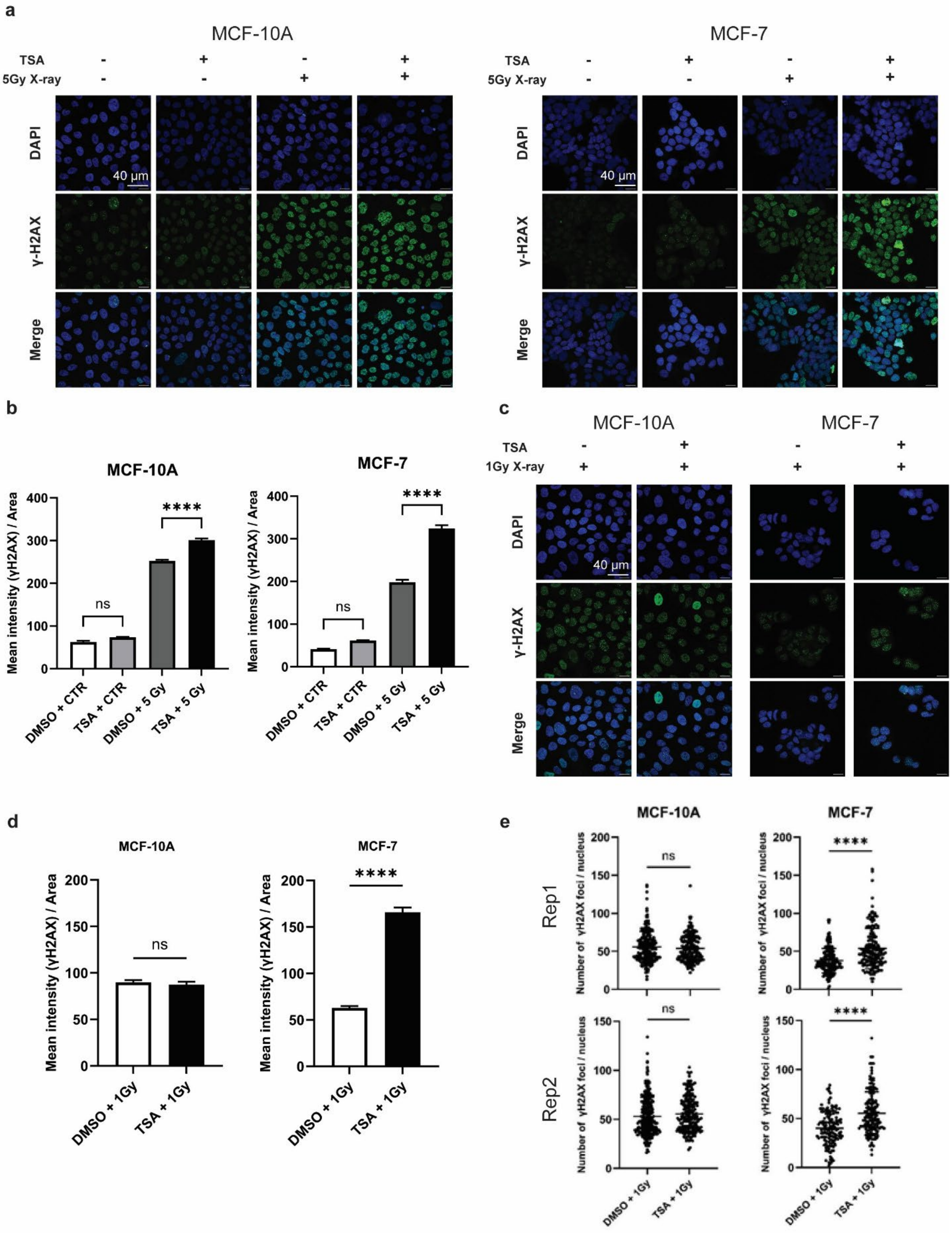


**Supplementary Figure 4. TSA enhances X-ray induced DNA damage in healthy and cancer cells differently for different radiation doses – biological replicate experiment.** This figure presents data from independent biological replicates of the experiments shown in Figures 2 and 3. (**a**)

MCF-10A (left) and MCF-7 (right) cells were pretreated with or without 500 nM TSA, irradiated with 5 Gy X-rays, and stained with DAPI (Blue) and anti-γH2AX (Green) 30 minutes after radiation. Scale bar: 40 µm (shown in top right image, same for all panels). (**b**) Quantification of average γH2AX intensity per nucleus area as in (**a**) Bars show mean +/- SEM averaged across all measured nuclei (5 fields imaged per condition, N=250 – 350 nuclei per condition) (** $p<0.01$, **** $p < 0.0001$, one-way ANOVA.). (**c**) MCF-10A (left) and MCF-7 (right) cells were pretreated with or without 500 nM TSA, irradiated with 1 Gy X-rays, and stained with DAPI (Blue) and anti-γH2AX (Green) 30 minutes after radiation. Scale bar: 40 µm (shown in top right image, same for all panels) (**d**) Quantification of average γH2AX intensity per nucleus area as in (**c**) Bars show mean +/- SEM averaged across all measured nuclei (5 fields imaged per condition, N=150 – 300 nuclei per condition) (**** $p < 0.0001$, one-way ANOVA.). **(e)** Quantification of γH2AX foci per nucleus for the 1 Gy irradiation data shown in panel (c). 5 fields imaged per condition, N=150 – 300 nuclei per condition.  Points are plotted for each nucleus and the mean is indicated as a horizontal line (**** $p < 0.0001$, one-way ANOVA.).

**a**

**A375 gene expression after 2 hr 0.5 uM TSA treatment**

*(Data from Vinayak et al., Nat Comm 2025)*

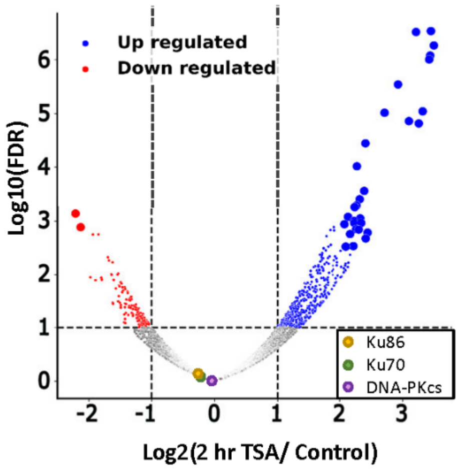


**b**

**Gene Ontology Enrichment for Differentially Expressed Genes for A375 + 2h TSA (>1.5 log2FC)**

Upregulated genes (240)

| GO-term | Description | # Genes | FDR |
|---|---|---|---|
| GO:0010632 | Regulation of epithelial cell migration | 13 | 0.0076 |
| GO:0010959 | Regulation of metal ion transport | 17 | 0.0086 |
| GO:0010646 | Regulation of cell communication | 70 | 0.0044 |
| GO:0045595 | Regulation of cell differentiation | 38 | 0.0135 |
| GO:0051716 | Cellular response to stimulus | 108 | 0.0076 |

Downregulated genes (49)

| GO-term | Description | # Genes | FDR |
|---|---|---|---|
| GO:0019219 | Regulation of nucleobase-containing compound metabolic process | 26 | 0.0095 |
| GO:0140110 | Transcription regulator activity | 15 | 0.0209 |
| GO:0043565 | Sequence-specific DNA binding | 16 | 0.0078 |

**c**

**MCF7 100 uM 6 h TSA treatment, vs. DMSO Gene Expression** *(GSE252117)*

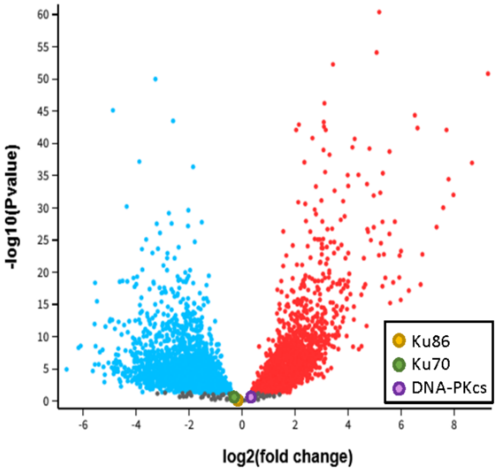


**d**

**Gene Ontology Enrichment for MCF7 + 6h TSA Differentially Expressed Genes (>1.5 log2FC)**

Upregulated genes (489)

| GO-term | Description | # Genes | FDR |
|---|---|---|---|
| GO:0140352 | Export from cell | 31 | 0.0216 |
| GO:0032940 | Secretion by cell | 28 | 0.0224 |
| GO:0008092 | Cytoskeletal protein binding | 55 | 0.00038 |
| GO:0042995 | Cell projection | 107 | 1.48e-06 |

Downregulated genes (380)

| GO-term | Description | # Genes | FDR |
|---|---|---|---|
| GO:0010468 | Regulation of gene expression | 215 | 2.41e-35 |
| GO:0051276 | Chromosome organization | 50 | 1.98e-07 |
| GO:0009725 | Response to hormone | 42 | 8.75e-07 |
| GO:0016570 | Histone modification | 27 | 1.18e-05 |
| GO:0045595 | Regulation of cell differentiation | 62 | 3.30e-05 |
| GO:0060548 | Negative regulation of cell death | 43 | 0.00042 |
| GO:0080135 | Regulation of cellular response to stress | 32 | 0.0025 |
| GO:0006974 | Cellular response to DNA damage stimulus | 32 | 0.0051 |
| GO:0016573 | Histone acetylation | 11 | 0.0094 |

**e**

**DNA damage response genes downregulated by 6 h TSA, MCF-7**

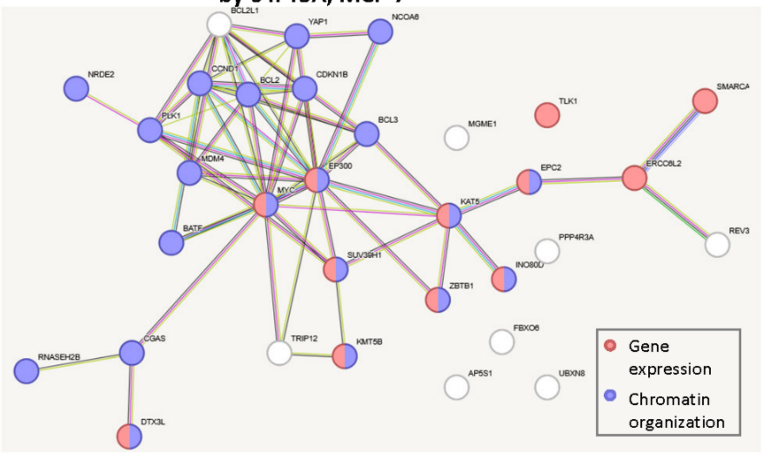

**Supplementary Figure 5. Transient TSA treatment has minimal effect on DNA repair gene expression in A375 and MCF7 cells.** (**a**) Differential expression analysis reveals genes up and downregulated by 2 hr, 500 nM TSA treatment in A375 cells, but shows that NHEJ pathway components Ku86, Ku70, and DNA-PKcs (colored circles on plot) are not affected. Figure adapted from Vinayak et al., 2025 used with permission of the Creative Commons Attribution 4.0 International License. (**b**) Gene ontology enrichment analysis of the differentially expressed genes (adjusted $p<0.05$, log2FC>1.5) from (a) performed in STRING and non-redundant categories selected for display. The number of genes in each functional category is shown in the table along with the false discovery rate adjusted enrichment significance. (**c**) Differential gene expression analysis was performed with GEO2R comparing 6 h treatment of MCF7 cells with TSA to control cells, using data published in GSE252117. NHEJ pathway components Ku86, Ku70, and DNA-PKcs (colored circles on plot) are not affected. (**d**) Gene ontology enrichment analysis of the differentially expressed genes (adjusted $p<0.05$, log2FC>1.5) from (c) performed in STRING and non-redundant categories selected for display. The number of genes in each functional category is shown in the table along with the false discovery rate adjusted enrichment significance. (**e**) STRING was used to create an interaction network for all genes in the Cellular response to DNA damage stimulus category found to be downregulated in MCF7 cells by TSA treatment in (d). Most of the proteins are found in the functional categories of gene expression regulation (red nodes) or chromatin organization (blue nodes).

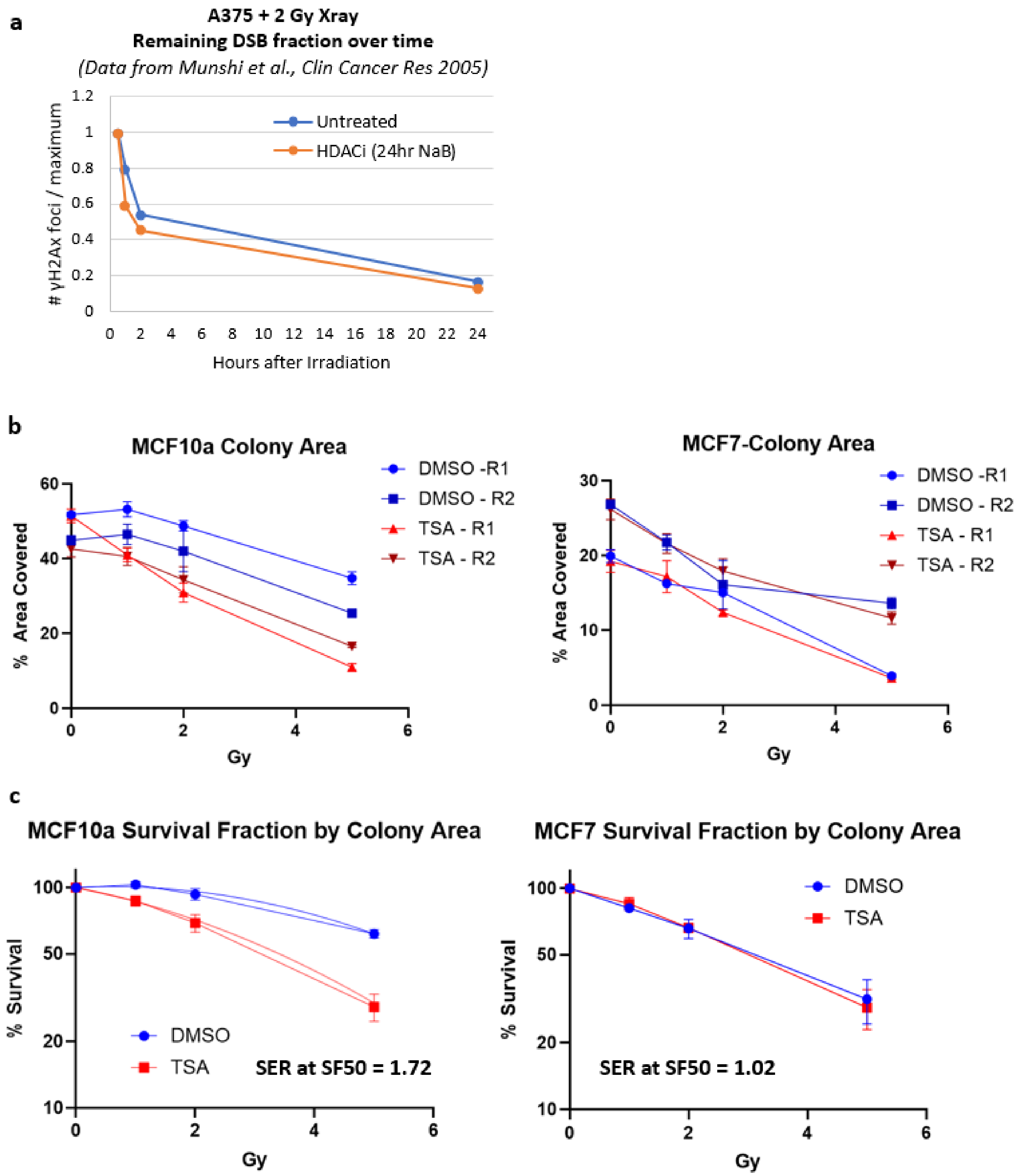


**Supplementary Figure 6. Alternative quantification of long term effects of HDACi + X-ray.** (**a**) Data for γH2Ax foci counts were collected from Munshi et al. 2005 for A375 cells pre-treated with 24 h HDACi vs. no treatment and then 2 Gy X-ray irradiation. DNA damage foci over time were normalized to the maximum value at the first timepoint, enabling comparison of repair rate on the same scale between the two conditions. No evident difference in repair rate over time is observed. (**b**) Clonogenic assay plate images (from Figure 4) were quantified using the

Colony Area approach (ImageJ plugin). Examining the un-normalized colony area measurements in each replicate shows that TSA has no effect on un-irradiated colony growth. (**c**) Plotting the data from (b) as a % survival curve (normalizing the data from each replicate to its un-irradiated starting point and plotting on a log scale) shows a modest radiosensitizing effect of TSA on MCF10a cells and a less notable effect by this quantification method in MCF7 cells. The sensitizer enhancement ratio (SER) at survival fraction 50% was calculated from a linear-quadratic fit (curve fit shown on MCF-10A plot).

**a**

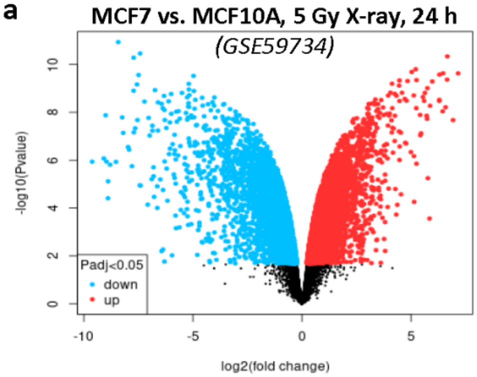


**b**

**Functional Enrichment for 5 Gy MCF7 vs. 10A Differentially Expressed Genes (>1.5 log2FC)**

Higher in MCF7 (725)

| Pathway | Description | # Genes | FDR |
|---|---|---|---|
| hsa03430 | Mismatch repair | 8 | 0.00047 |
| hsa03030 | DNA replication | 12 | 2.4E-05 |
| hsa03440 | Homologous recombination | 11 | 0.00032 |
| hsa03083 | Polycomb repressive complex | 19 | 1.1E-05 |
| hsa03410 | Base excision repair | 10 | 0.00177 |
| hsa03013 | Nucleocytoplasmic transport | 22 | 1.1E-05 |
| hsa01524 | Platinum drug resistance | 15 | 0.00035 |
| hsa01522 | Endocrine resistance | 17 | 0.00046 |
| hsa05235 | PD-L1 expression and PD-1 checkpoint pathway in cancer | 15 | 0.00177 |
| hsa04919 | Thyroid hormone signaling pathway | 20 | 0.0003 |
| hsa04915 | Estrogen signaling pathway | 22 | 0.0003 |
| hsa04110 | Cell cycle | 25 | 0.0001 |
| hsa04218 | Cellular senescence | 21 | 0.00188 |

Higher in MCF-10A (843)

| GO-term | Description | # Genes | FDR |
|---|---|---|---|
| GO:0042060 | Wound healing | 56 | 5.11E-15 |
| GO:0030155 | Regulation of cell adhesion | 99 | 5.06E-19 |
| GO:0030334 | Regulation of cell migration | 109 | 5.06E-19 |
| GO:0001817 | Regulation of cytokine production | 90 | 1.65E-16 |
| GO:0071219 | Regulation of MAPK cascade | 80 | 2.26E-14 |
| GO:0033993 | Response to lipid | 89 | 1.38E-13 |

**c**

**Notable genes differently regulated in both 5 Gy and 9 Gy radiation in MCF7 vs. MCF10A**

| | | MCF7 vs. 10A | | | |
|---|---|---|---|---|---|
| | | **5 Gy** *(GSE59734)* | | **9 Gy** *(GSE108962)* | |
| **function** | **gene** | **logFC** | **adjPval** | **logFC** | **adjPval** |
| cell death | CASP8 | -1.28 | 0.0118 | -0.98 | 1E-05 |
| cell death | IFI6 | 2.21 | 0.006 | 1.37 | 7E-07 |
| cell migration | CXCL3 | -6.28 | 4E-06 | -1.10 | 0.0085 |
| cell migration | C3AR1 | -3.57 | 0.0041 | -1.24 | 0.0003 |
| cell migration | PTPN14 | -1.19 | 0.0052 | -1.01 | 2E-05 |
| chromatin | HDAC7 | -1.08 | 0.0015 | -1.39 | 9E-05 |
| chromatin | JUND | 0.83 | 0.0109 | 1.54 | 1E-09 |
| chromatin | TFAP2C | 3.14 | 4E-06 | 1.52 | 0.0002 |
| chromatin | DNMT3A | 2.82 | 8E-06 | 2.60 | 0.006 |
| chromatin | MGMT | 1.10 | 0.0001 | 0.66 | 0.0015 |
| chromatin | CENPA | 1.52 | 0.0152 | 0.91 | 0.0027 |
| chromatin | CITED2 | 2.71 | 0.0002 | 0.99 | 7E-07 |
| DNA repair | XRCC1 | 1.57 | 0.0001 | 1.23 | 5E-06 |
| ECM | SERPINH1 | -1.18 | 1E-04 | -1.27 | 0.0003 |
| ECM | COL6A1 | -1.15 | 0.0024 | -1.79 | 2E-06 |

**Supplementary Figure 7. Differential gene regulation in response to X-ray in MCF7 vs. MCF10a cells.** (**a**) Differential gene expression analysis was performed using GEO2R to compare MCF7 and MCF710A transcriptomes 24 h after exposure to 5 Gy X-ray, using data published in GSE59734. (**b**) Gene ontology enrichment analysis of the differentially expressed genes (adjusted p<0.05, log2FC>1.5) from (a) was performed using ShinyGO KEGG pathways and Gene Ontology terms. Non-redundant categories selected for display show extensive upregulation of DNA repair genes in MCF7 as compared to MCF10a after this X-ray treatment. The number of genes in each functional category is shown in the table along with the false discovery rate adjusted enrichment significance. (**c**) Differential expression analysis was performed for MCF7 vs. MCF10A after 9 Gy exposure and then genes that change in the same direction in 5 Gy and 9 Gy datasets were considered. Genes from biologically relevant functional categories are highlighted here along with log2FC values and FDR adjusted P-values.
